# The sleep-wake distribution contributes to the peripheral rhythms in PERIOD-2

**DOI:** 10.1101/2020.07.25.221101

**Authors:** Marieke M.B. Hoekstra, Maxime Jan, Yann Emmenegger, Paul Franken

**Affiliations:** Center for Integrative Genomics, University of Lausanne, Lausanne, Switzerland; UK Dementia Research Institute at Imperial College London, Division of Brain Sciences, London, United Kingdom

**Keywords:** circadian rhythm, sleep, waking, *Period-2*, bioluminescence, locomotor activity, modeling, kidney, cortex

## Abstract

In the mouse, *Period-2* (*Per2*) expression in tissues peripheral to the suprachiasmatic nuclei (SCN) increases during sleep deprivation and at times of the day when animals are predominantly awake spontaneously, suggesting that the circadian sleep-wake distribution directly contributes to the daily rhythms in *Per2*. We found support for this hypothesis by recording sleep-wake state alongside PER2 bioluminescence in freely behaving mice, demonstrating that PER2 increases during spontaneous waking and decreases during sleep. The temporary reinstatement of PER2 rhythmicity in behaviorally arrhythmic SCN-lesioned mice submitted to daily recurring sleep deprivations substantiates our hypothesis. Mathematical modelling revealed that PER2 dynamics can be described by a damped harmonic oscillator driven by two forces: a sleep-wake-dependent force and a SCN-independent circadian force. Our work underscores the notion that in peripheral tissues the clock gene circuitry integrates sleep-wake information and could thereby contribute to behavioral adaptability to respond to homeostatic requirements.

## Introduction

The sleep-wake distribution is coordinated by the interaction of a circadian and a homeostatic process (Daan et al., 1984). The biological substrates underlying the circadian process are relatively well understood: circadian rhythms in overt behavior of mammals are generated by the suprachiasmatic nuclei (SCN) located in the hypothalamus (Hastings et al., 2018). At the molecular level, so called ‘clock genes’ interact through negative transcriptional/translational feedback loops (TTFLs), where the CLOCK/NPAS2:ARNTL (BMAL1) heterodimers drive the transcription of their target genes, among them the *Period* (*Per1,2*) and *Cryptochrome* (*Cry1,2*) genes. Subsequently, PER and CRY proteins assemble into repressor complexes that inhibit CLOCK/NPAS2:ARNTL-mediated transcription, including their own. The resulting reduction of the repressor complex allows a new cycle to start. This feedback loop, present in almost each cell of the mammalian body, interacts with other molecular pathways, together ensuring a period of ~24h (Hastings et al., 2018). The SCN synchronizes peripheral clock gene expression rhythms through its rhythmic behavioral, electrical, and humoral output generated across the day (Schibler et al., 2015).

Accumulating evidence suggests that, perhaps surprisingly, clock genes are also involved in the homeostatic aspect of sleep regulation (Franken, 2013). This is illustrated by the sleep-deprivation (SD) induced increase in the expression of the clock gene *Per2* in tissues peripheral to the SCN, including the cerebral cortex, liver, and kidney (Curie et al., 2013, 2015; Franken et al., 2007; Maret et al., 2007). Moreover, the highest level of peripheral *Per2* expression is reached after the time of day mice were awake most, suggesting that also during spontaneous periods of waking *Per2* expression accumulates. Accordingly, lesioning of the SCN, which eliminates the circadian sleep-wake distribution, attenuates the circadian amplitude of clock gene transcripts and proteins in peripheral tissues (Akhtar et al., 2002; Curie et al., 2015; Tahara et al., 2012; Sinturel et al., 2021). Together, these studies suggest that sleeping and waking are important contributors to clock gene expression but dissecting the contribution of the sleep-wake distribution and circadian time is challenging, because the two change in parallel.

By simultaneously recording electroencephalogram (EEG), electromyogram (EMG), locomotor activity and PER2-dependent bioluminescence signals from cortex and kidney in freely behaving mice, we established that the circadian sleep-wake distribution importantly contributes to the daily rhythmic changes in central and peripheral PER2 levels. To further test this hypothesis, we predicted that i) in behaviorally arrhythmic SCN-lesioned (SCNx) mice, daily recurring SDs mimicking a circadian sleep-wake distribution, will temporarily reinstate high amplitude PER2 bioluminescence rhythms and ii) in intact rhythmic animals, reducing the amplitude of the circadian sleep-wake distribution will result in a reduced amplitude of PER2 rhythms. While daily SDs indeed enhanced the amplitude of peripheral PER2 rhythms in SCNx mice, the protocol used to reduce the amplitude of the sleep-wake distribution did not reduce PER2 amplitude in all mice. To reconcile the sleep-wake driven and circadian aspects of PER2 dynamics, we implemented a mathematical model in which waking represents a force that sets in motion a harmonic oscillator describing PER2 dynamics, and found that the sleep-wake distribution, also under undisturbed conditions, is an important contributor to the daily changes in PER2 bioluminescence. Moreover, we discovered a second, SCN and sleep-wake independent force with a circadian period that underlay the residual circadian PER2 rhythms in SCNx mice, and that the phase relationship between these two forces is important for predicting the amplitude response in PER2 rhythms to sleep-wake perturbations.

## Results

To quantify PER2 levels, we used mice expressing a knock-in (KI) construct encoding a fused PER2::LUCIFERASE (PER2::LUC) protein and in which changes in emitted bioluminescence can be used as proxy for changes in PER2 protein levels (Yoo et al., 2004). *Per2::Luc* KI mice have been used to follow clock gene expression *in vivo* (Curie et al., 2015; Ohnishi et al., 2014; Tahara et al., 2012; van der Vinne et al., 2018). However, in these studies mice had to be anaesthetized for each measurement, while in the set-up used in our study [RT-Biolumicorder (C. Saini et al., 2013)], we assessed PER2-bioluminenscence continuously in freely moving mice, in central and peripheral tissues. For the central quantification of bioluminescence, we used mice in which the PER2::LUC construct was back-crossed onto a C57BL/6J (B6) background (see Methods). For the experiments that quantified bioluminescence in the periphery, we used hairless SKH1 mice carrying the *Per2::Luc* KI construct, because lack of fur allows for the unobstructed measurement of emitted photons (see also Suppl. Fig. 1). Under standard LD12:12 conditions, SKH1 mice exhibited sleep-wake patterns characteristic of mice, i.e. during the light phase they spent more time in both Non-Rapid-Eye-Movement (NREM) sleep and REM sleep relative to the dark phase, the latter being their habitual active phase. Moreover, they showed the typical sleep homeostatic response to a 6h SD during the first 18h of recovery, both in sleep time and EEG delta power, although the increase in REM sleep did not reach significance levels (Suppl. Fig. 2).

In two pilot experiments we optimized our experimental set-up. We established that the most important source contributing to the peripheral bioluminescence signals in the SKH1 mice are the kidneys (Suppl. Fig. 3A). We have previously shown that the central bioluminescence signal obtained in B6 mice is of cortical origin (Curie et al. 2015). By using mice constitutively expressing luciferase (Y.-A. Cao et al., 2004), we determined that its substrate luciferin is best delivered through implantable osmotic mini-pumps compared to administration through the drinking water. Under the latter condition, strong daily rhythms in bioluminescence were observed, likely as a result of rhythms in drinking behavior, thereby driving luciferin availability (Suppl. Fig. 3B). To record peripheral bioluminescence, we implanted mice subcutaneously with mini-pumps in a short (5min) intervention. To record central bioluminescence, luciferin was administered centrally through a cannula connected to a subcutaneously placed mini-pump, requiring stereotactic surgery. In addition, the skull was thinned and equipped with a glass cone facilitating passage of photons through the skull (Curie et al. 2015; see Methods for details).

### SLEEP-WAKE STATE AFFECTS PER2 BIOLUMINESCENCE

It now has been well documented that enforced wakefulness affects *Per2* mRNA and protein levels in various tissues and mammalian species (Franken, 2013; Hoekstra et al., 2019; Möller-Levet et al., 2013; Vassalli & Franken, 2017), but it is not known whether circadian rhythms in spontaneous sleep-wake behavior contribute to the daily changes in PER2 levels. To address this question, we equipped *Per2::Luc* KI B6 and SKH1 mice (n=6 and 5, respectively) with wireless EEG/EMG recorders and monitored simultaneously sleep-wake state, PER2 bioluminescence and locomotor activity under constant darkness (DD; see Suppl. Fig. 1). With this set-up, we reproduced the circadian changes in PER2 bioluminescence as well as the response to a 6h SD (Fig. 1A) as described previously (Curie et al., 2015) (see material and methods for explanation of the SD procedure). Similar to what was observed in that publication, SD elicited a tissue-specific response, with an immediate increase in central PER2-bioluminescence after SD, while in the periphery the response was delayed and PER2 bioluminescence increases were observed in the 2^nd^ and 5^th^ hour of recovery (Fig. 1A). In addition to these direct effects on PER2 bioluminescence within the 1st 6h of recovery, the SD also caused a long-term reduction of rhythm amplitude during both recovery days; i.e. the first (REC1) and last (REC2) 24h period of recovery. In the periphery this decrease amounted to ca. 30% on both days [REC1: 17.5-41.1%; REC2 18.7-42.4%, 95% confidence intervals (95%-CI)]. In the central recordings PER2-bioluminescence amplitude decreased significantly during REC1 [32% (10.2-54.5%)], whereas the decrease during REC2 [22% (0.06-44.8%)] no longer reached significance levels [linear mixed model with fixed conditional effect (‘BSL’, ‘REC1’, ‘REC2’) and random intercept effect (‘Mouse’); Periphery: BSL vs. REC1 p=0.0014; vs. REC2 p=0.0011; Central: BSL vs. REC1 p=0.022; vs. REC2 p=0.082; sinewave fitted baseline amplitudes: periphery 0.258 (0.15-0.36); central 0.193 (0.08-0.31 a.u.)]. This long-term reduction of PER2-biolumnescence is reminiscent of the long-term SD effects on rhythm amplitude we observed for *Per2* expression and for other clock genes in the cortex (Hor et al. 2019).

**Figure 1:**
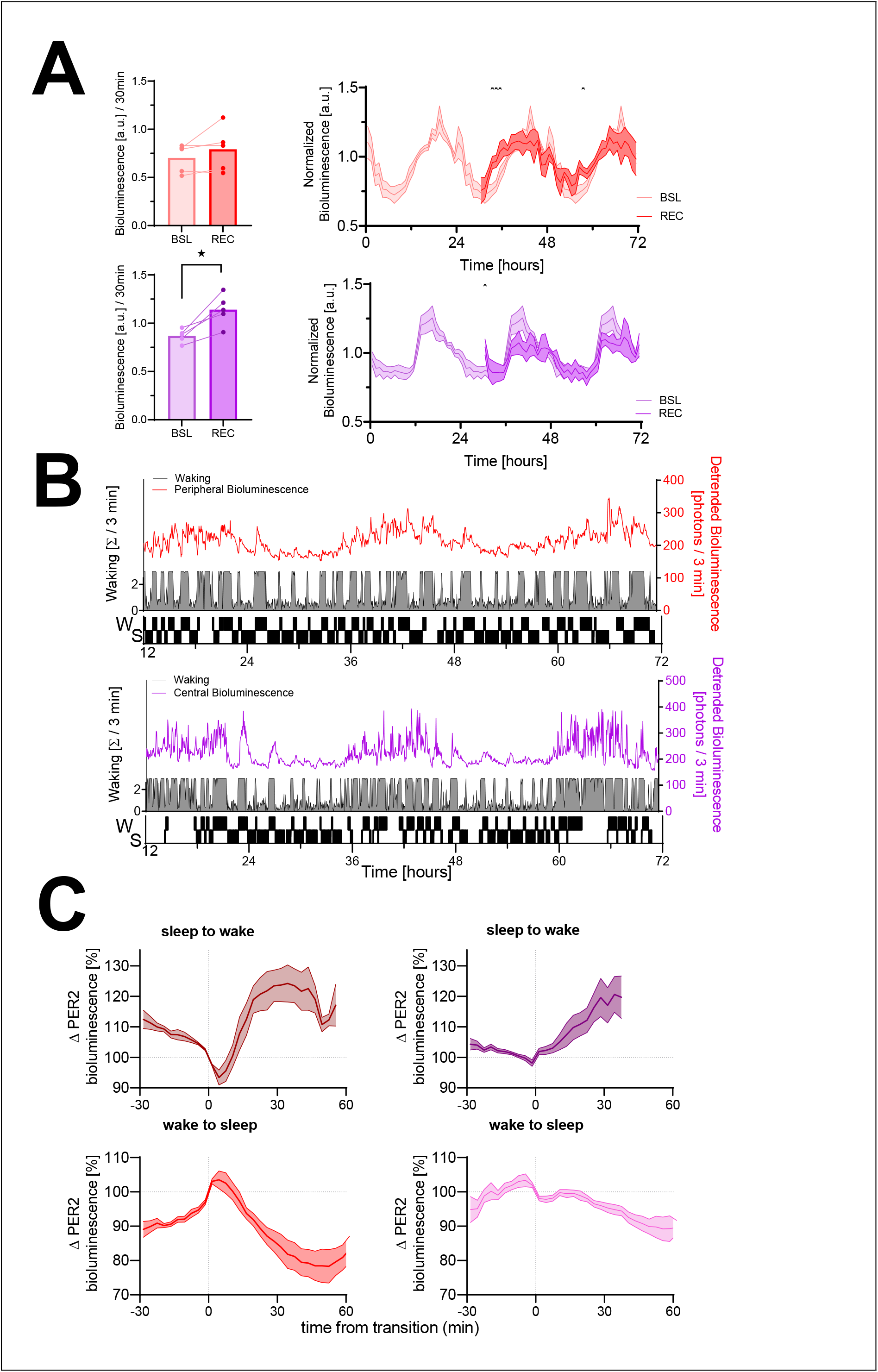
The sleep-wake distribution contributes to changes in PER2 bioluminescence. Red = peripheral bioluminescence, purple = central bioluminescence. Left panels: PER2 bioluminescence measured in the first 30min of recovery (REC) after sleep deprivation compared to levels reached at this circadian time during baseline (BSL). Sleep deprivation elicited an acute response in central PER2 (lower bar graph: t(4)=4.4, p=0.012), but not in the periphery (upper bar graph, t(4)=1.5, p=0.21). Right panels: Under baseline conditions (lighter graphs, time 0-24h repeated three times, average of 48h baseline), PER2 bioluminescence showed a circadian rhythm both in the periphery (left) and central (right). Sleep deprivation from ZT0 to −6 (times under preceding LD conditions), affected PER2 bioluminescence during recovery (2-way rANOVA w/ Condition x Time, periphery: F(41,164)=2.1, p=0.0007; central: F(41,164)=1.8, p=0.004). Asterisks indicate significant differences assessed by post-hoc paired t-tests. Bioluminescence is expressed as a fraction of the individual average bioluminescence during the experiment and depicted in 1h intervals as mean±SEM for 5 mice (peripheral and central). (**B**) A sleep-wake state recording (grey area plot represents wakefulness in consecutive 3-min intervals) combined with peripheral (red line, upper graph) and central PER2 bioluminescence (purple line, lower graph) in two mice during baseline. Note that besides the circadian oscillation in PER2 bioluminescence, there are marked increases and decreases in PER2 bioluminescence. The hypnogram (lower part of the graph) illustrates that the rapidly evoked changes in PER2 bioluminescence are related to periods of sleeping (S) and waking (W). This hypnogram is discontinuous as it depicts only the SW transitions selected in this mouse for the analysis in C. (**C**) Changes in PER2 bioluminescence associated with transitions from sleep to wake (top) and wake to sleep (bottom) of peripheral (left, n=5) and cortical (right, n=6) origin.

Although a circadian modulation of PER2 bioluminescence of both peripheral and central origin is evident (see Fig. 1B and Suppl. Fig. 4), we observed additional changes in bioluminescence that occurred simultaneously with changes in sleep-wake state, indicating that PER2 bioluminescence increases during wake-dominated periods and decreases during sleep-dominated periods in both tissues. To quantify this observation, transitions from sleep (irrespective of sleep state) to wake and from wake to sleep were selected (see Material and Methods for selection criteria). Examples of the selected transitions are indicated in Fig. 1B as a hypnogram. A similar number of sleep-to-wake and wake-to-sleep transitions passed selection criteria during the two-and-a-half baseline days (periphery: 31.2±3.9 and 29.0±3.9; central: 25.6±1.4 and 24.6±1.4, respectively; mean±SEM). Transitions obtained in mice in which central bioluminescence was recorded were shorter (longest common wake period after sleep-to-wake transitions: 39 vs. 57min; longest common sleep period after wake-to-sleep transitions: 63 vs. 75min, for central and periphery, respectively). Although PER2-bioluminescence increased during wakefulness and decreased during sleep in both tissues, tissue-specific differences were observed (Fig. 1C). At sleep-to-wake transitions tissue differences concerned an initial decrease in peripheral PER2-bioluminescence after wake onset, followed by a steep increase saturating at 125%. In contrast, central levels of PER2 bioluminescence increased from the start and followed a linear time course throughout the waking period. Despite these different dynamics, similar bioluminescence levels were reached in both tissues. After wake-to-sleep transitions peripheral PER2 bioluminescence initially increased before quickly decreasing and then leveling out at around 84% (Fig. 1C). In contrast, the central signal decreased linearly throughout the sleep period reaching levels of 91% at the end.

Taken together, two observations indicate that the sleep-wake distribution importantly contributes to both central and peripheral PER2-bioluminescence dynamics: 1) enforcing wakefulness has both acute and long-term effects on PER2 bioluminescence, 2) the changes in PER2 bioluminescence at sleep-wake transitions demonstrate that also spontaneous waking is associated with an increase in PER2 bioluminescence and sleep with a decrease, and 3) that mice are spontaneously awake more when PER2 bioluminescence is high.

The central PER-bioluminescence signals tended to be noisier than peripheral signals [signal-to-noise ratio: −1.15±1.0 and 0.37±1.0 dB, respectively; estimated on 3-min values in baseline of the 6h-SD experiments according to (Leise et al., 2012)]. Moreover, the peripheral signal was easier to obtain and minimally invasive to the animal. We therefore decided for the next experiments, to focus on peripheral PER2 bioluminescence using the *Per2::Luc KI* SKH1 mice. Because waking correlates highly with LMA (Suppl. Fig. 5), we recorded LMA as proxy for wakefulness to avoid invasive EEG/EMG surgery and mice having to adapt to a ca. 2.7 g recorder mounted on the head.

### MODULATING THE AMPLITUDE OF THE SLEEP-WAKE DISTRIBUTION AND ITS INFLUENCE ON PER2 BIOLUMINESCENCE

The SCN is the main driver of the circadian sleep-wake distribution because animals in which the SCN is lesioned (SCNx) lack a circadian organization of sleep-wake behavior under constant conditions (Baker et al., 2005; Edgar et al., 1993). Under these conditions, the amplitude of clock gene expression in peripheral organs is significantly reduced, but not eliminated (Akhtar et al., 2002; Curie et al., 2015; Tahara et al., 2012; Sinturel et al., 2021). Given our results in freely behaving mice, we expect that imposing a rhythmic sleep-wake distribution in SCNx mice will reinstate high-amplitude rhythmicity in PER2 bioluminescence. Conversely, reducing the amplitude of the circadian sleep-wake distribution in SCN-intact mice is expected to reduce the amplitude of PER2 bioluminescence. We tested these predictions in two complementary experiments (Suppl. Fig. 1B and -C). In the first experiment, we enforced a daily sleep-wake rhythm in arrhythmic SCNx mice by sleep depriving them for 4h at 24h intervals during 4 subsequent days. In the second experiment, we aimed to acutely reduce the circadian distribution of sleeping and waking in intact mice according to a ‘2hOnOff’ protocol, comprising of twelve 2h SDs each followed by a 2h sleep opportunity ‘window’ (SOW), previously utilized in the rat (Yasenkov & Deboer, 2010).

#### Repeated sleep deprivations in SCNx mice temporarily reinstate a circadian rhythm in PER2 bioluminescence

Lesioning the SCN rendered locomotor activity arrhythmic under DD conditions (Suppl. Fig. 6). In arrhythmic SCNx mice, we confirmed that rhythms in PER2 bioluminescence were strongly reduced, but not completely abolished (Fig. 2A, -B, and -C). During the second baseline measurement after SCNx (BSL2 in Fig. 2B), we observed in 2 of the 4 mice erratic high values which we cannot readily explain especially because in the subsequent four SD days the variance in the PER2 bioluminescence signal among mice was again as small as in the earlier recordings (Fig. 2A vs. -B). The erratic values in these 2 mice led to a >10-fold increase in the residual sum of squares, indicating a poorer fit, and to a higher amplitude of the fitted sinewaves in the second compared to the first baseline recording (Fig. 2C, BSL1 vs. BSL2). The repeated SDs induced a robust oscillation in the PER2 bioluminescence signal, restoring amplitude to the levels observed prior to SCN lesioning and higher than those observed in the baseline recordings (Fig. 2C). The latter observation shows that the increased amplitude during the SD did not result from a SD-mediated alignment of different individual phases in baseline. Importantly, the oscillation continued after the end of the SDs while decreasing in amplitude (Fig. 2C).

**Figure 2:**
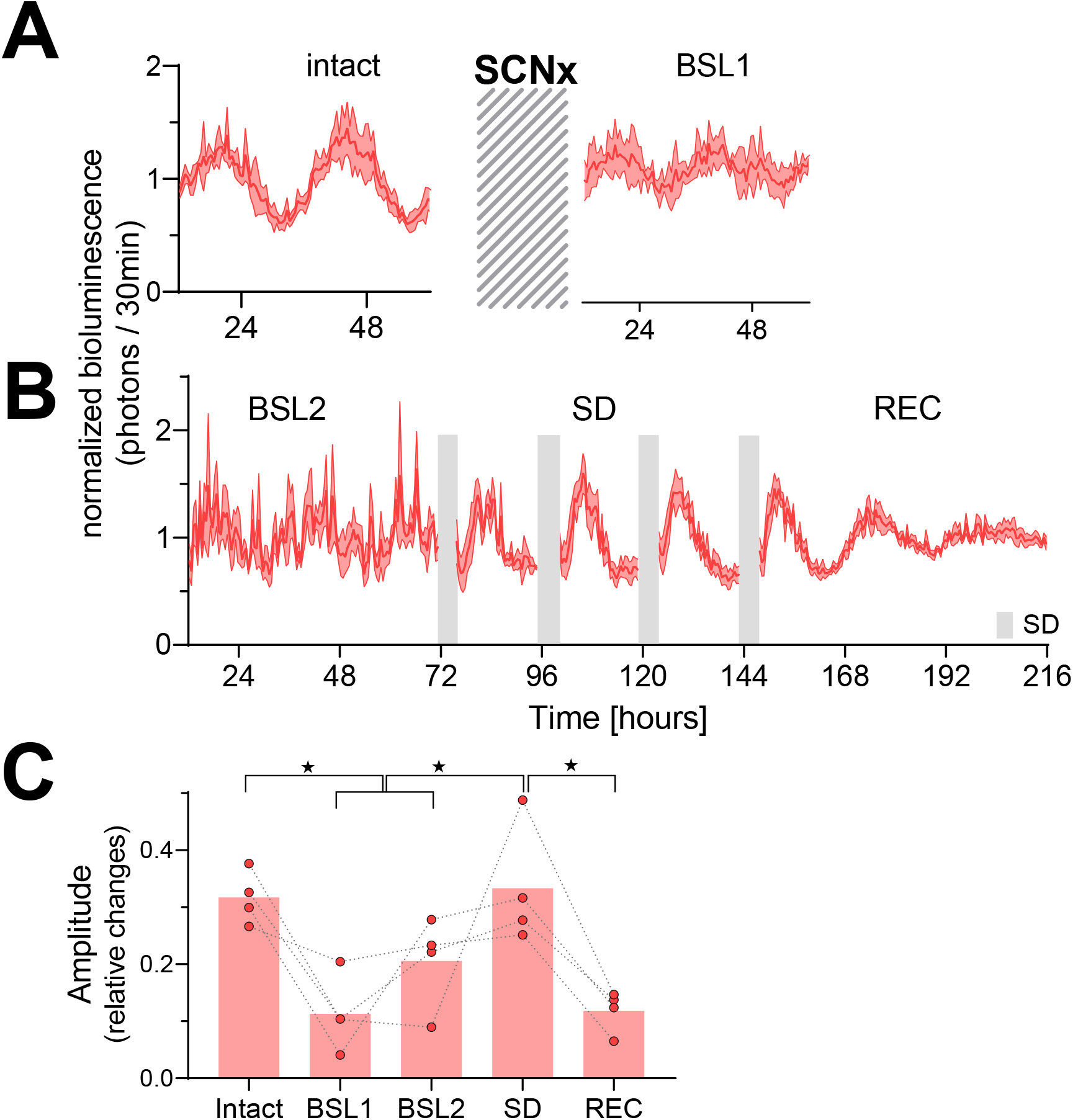
Temporary reinstatement of circadian PER2 oscillations in SCNx mice by repeated sleep deprivations. (**A**) Time course of PER2 bioluminescence across 2.5 days under baseline conditions (left) and after the SCN lesion (SCNx, right). Cross-hatched area represents approximately 5-week interval separating the 2 recordings during which the success of SCNx on locomotor activity was verified under DD (Suppl. Fig. 6). (**C**) Four 4h sleep deprivations (SD, grey bars) repeated daily at the beginning of the light phase under the preceding LD conditions, reinstate rhythmic PER2 bioluminescence (n=4). Abbreviation of experimental conditions: “intact”: baseline prior to SCNx; “BSL1”: baseline after DD locomotor activity recordings; “BSL2”: baseline immediately preceding the SDs; “SD”: 20h recordings between the four 4h SDs; “REC”: recovery after the four SDs. Data are depicted as mean±SEM (n=4). (**C**) Effect of SCNx and sleep deprivation on PER2 bioluminescence amplitude as determined by sinewave fitting [linear mixed model with fixed conditional effects (‘Intact’, ‘BSL’, ‘SD’ and ‘REC’) and random intercept effect (‘Mouse’) followed by Tukey’s post-hoc tests; intact vs. BSL (BSL1 and BSL2), p=0.0045; intact vs. REC, p=0.0015; SD vs. BSL: p=0.0013; SD vs. REC: p<0.001; ★: p<0.05].

#### The 2hOnOff protocol reduced the amplitude of the circadian sleep-wake distribution without impacting the circadian expression of PER2

In the next experiment, we aimed at reducing the amplitude of the circadian sleep-wake distribution in intact mice using the 2-day 2hOnOff protocol. We measured sleep-wake state and PER2 bioluminescence during baseline conditions, the 2hOnOff procedure and the two recovery days in two cohorts of mice. In the first cohort, PER2 bioluminescence was monitored, and mice were taken out of the RT-Biolumicorder for the 2h SDs. Bioluminescence was therefore quantified only during the SOWs, thus biasing the read-out of PER2 bioluminescence during the 2hOnOff protocol towards levels reached during sleep. A second cohort was implanted with tethered EEG/EMG electrodes to determine the efficacy of the 2hOnOff protocol in reducing the sleep-wake distribution amplitude.

As expected, mice exhibited a circadian PER2-bioluminescence rhythm under baseline conditions (Fig. 3A). Contrary to expectation, the 2hOnOff protocol did not significantly decrease the ongoing circadian PER2 oscillation when analyzed at the group level (Fig. 3B). When inspecting the individual responses, 4 out of the 6 mice did show a consistent 27% reduction (range 23-31%) in PER2-biolumininescence amplitude during the 2-day 2hOnOff protocol, while in the remaining two mice amplitude increased by 40% (Fig. 3C). In all 6 mice, PER2 bioluminescence amplitude reverted to baseline values during recovery, irrespective of whether it was increased or decreased during the 2hOnOff protocol. The distinct bimodal, opposing response in amplitude observed among individual mice and the subsequent reverting back to baseline suggests that the 2hOnOff protocol did affect the ongoing PER2 bioluminescence rhythm. This phenomenon might relate to factors not accounted for in our experimental design. We will address one potential contributing factor in the modeling section below.

**Figure 3:**
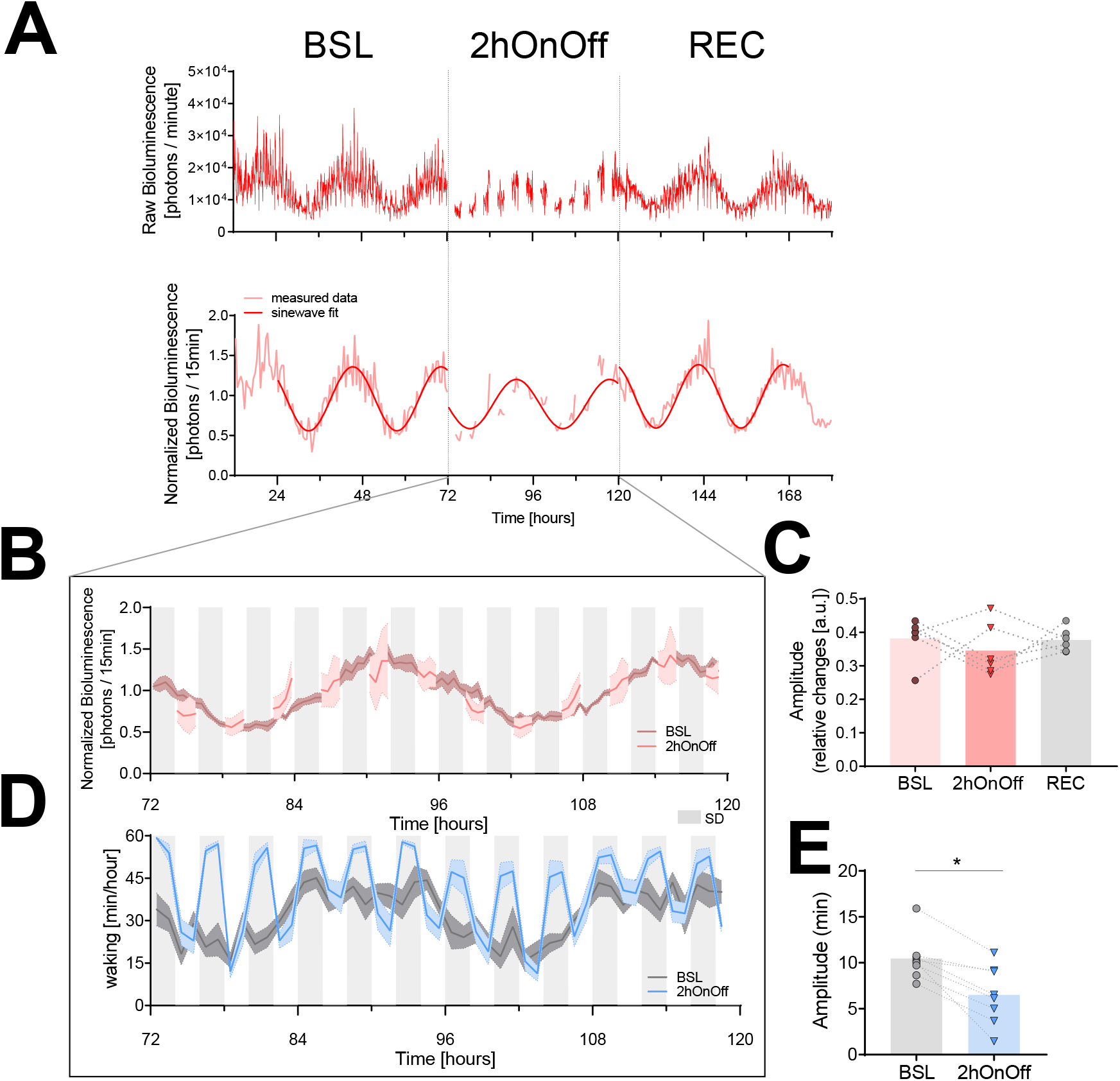
The 2hOnOff protocol reduced the circadian sleep-wake amplitude but did not consistently affect PER2 bioluminescence. (**A**) An example of a PER2-bioluminescence recording under baseline, 2hOnOff, and recovery conditions. Upper graph shows bioluminescence data as collected, whereas in the lower graph, the same data are linearly detrended, normalized (relative to the individual overall mean), and averaged over 30min intervals. A sinewave was fit to the data of the two baseline days, the two 2hOnOff days, and the two recovery days separately (see Materials and Methods). (**B**) The time course of PER2 bioluminescence under baseline conditions and during the 2hOnOff protocol (n=6, data depicted as mean ± SEM). Light grey squares below the graph mark the 2h SDs. (**C**) The amplitude of the PER2-bioluminescence rhythm decreased in 4 but increased in 2 mice resulting in an overall lack of an effect of the 2hOnOff protocol (paired t-test, t(5)=0.74, p=0.50; mean±SEM; BSL: 0.38±0.03; 2hOnOff: 0.35±:0.03; REC: 0.38±0.01). For all 6 animals the amplitude reverted to baseline during recovery. (**D**) The distribution of waking across the two baseline days (dark grey) and during the 2hOnOff protocol (blue) in EEG-implanted SKH1 mice (n=8, hourly values depicted as mean ± SEM) and a sinewave fit through the hourly average for visual comparison of BSL to SD. (**E**) Individual estimates of the amplitude of circadian changes in wakefulness during the 2hOnOff protocol were obtained by fitting a sine-wave function to the wakefulness present in consecutive 4h intervals (i.e. SD + SOW). This amplitude was smaller compared to the amplitude obtained in baseline using the same approach (paired t-test, t(7)=4.9, p=0.002; mean±SEM; BSL: 10.4±0.9; 2hOnOff: 6.5±1.1). However, a circadian modulation was still present under the 2hOnOff protocol (amplitude>0, 1-sample t-test, t(7)=5.8, p=0.0007). The sleep-wake distribution during recovery was not assessed. Note the overall higher levels of wakefulness during 2hOnOff compared to baseline.

We used the second cohort of mice to determine the efficacy with which the 2hOnOff protocol reduced the circadian sleep-wake distribution. In contrast to bioluminescence, the amplitude of the circadian sleep-wake distribution did decrease consistently in all mice (Fig. 3D and -E). The circadian rhythm in the sleep-wake distribution was, however, not eliminated because the sleep obtained during the 2h SDs as well as during the 2h SOWs both varied as a function of time of day (sleep during SDs: average: 12.4%, min-max: 4.9-23.0%; during SOWs: average: 54.3%, min-max: 35.7-76.4%). This was especially evident at the beginning of the subjective light phase of day 2, when the average time spent asleep during the SD reached 23%. The sleep obtained during the SDs at this time could be due to an substantial sleep pressure because of lost sleep during day 1 of the 2hOnOff protocol (total time spent asleep/day, mean±SEM; baseline: 11.1±0.1h; 2hOnOff: day 1: 7.3±0.3h, day 2: 8.6±0.7h, paired two-tailed t-test, BSL versus 2hOnOff-day 1: t(7)=8.86, p<0.001; BSL versus 2hOnOff-day2: t(7)=4.18, p=0.004), 2hOnOff-day1 versus -day2, t(7)=1.97, p=0.09), combined with the difficulty for the experimenters to visually detect and prevent sleep in pinkish hairless mice under dim-red light conditions.

### MODELING CIRCADIAN PER2 DYNAMICS

The results above demonstrate that both circadian and sleep-wake dependent factors need to be considered when studying PER2 dynamics. To understand how these two factors collectively generate the variance in PER2 bioluminescence, we put our experimental data into a theoretical framework and modeled changes in peripheral PER2 bioluminescence according to a driven harmonic oscillator (Curie et al., 2013). This type of oscillator is not self-sustained but depends on rhythmic forces to set in motion and maintain the oscillation. In the model, we assume the sleep-wake distribution to represent one such force. In the absence of rhythmic forces, the oscillator loses energy resulting in a gradual reduction of its amplitude according to the rate set by its damping constant (γ). Besides amplitude and timing (i.e. phase) of the recurring forces and the damping constant, the oscillatory system is further defined by a string constant 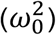 with the natural angular frequency (*ω*_0_) defining the intrinsic period. Because waking was not quantified alongside bioluminescence in the SCNx and the 2hOnOff experiments, we used locomotor activity (LMA) as a proxy for the driving force provided by wakefulness 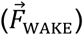. The 6h SD experiment showed that LMA and time spent awake are strongly correlated and that their hourly dynamics closely changed in parallel (Suppl. Fig. 5A and -B). Moreover, applying the model to this data set (see below) using either wakefulness or LMA as 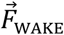 yielded similar fits (Suppl. Fig. 5C) demonstrating that LMA is an appropriate proxy for wakefulness for the purpose of our model. We performed the analyses at the group level; i.e., the mean LMA level of all mice was used to reflect 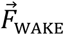, and the free parameters in the model were optimized by fitting the motion of the oscillator to the mean PER2-bioluminescence levels (see Material and Methods). Besides 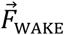, we implemented a circadian force to account for the residual PER2 bioluminescence rhythm observed in the SCNx mice (Fig. 2). We assumed that this additional, SCN and sleep-wake independent force reflects a peripheral circadian process 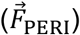 present in both intact and SCNx mice (Sinturel et al., 2021). This force was modeled as a sinewave with amplitude and phase as free parameters in both experiments while period was set to the period estimated from the baseline PER2-bioluminescence rhythm observed in the SCNx mice (23.7±0.6 h; n=4). A schematic overview of the influence of 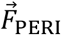 and 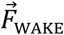 on PER2 bioluminescence is presented in Fig. 4A.

**Figure 4:**
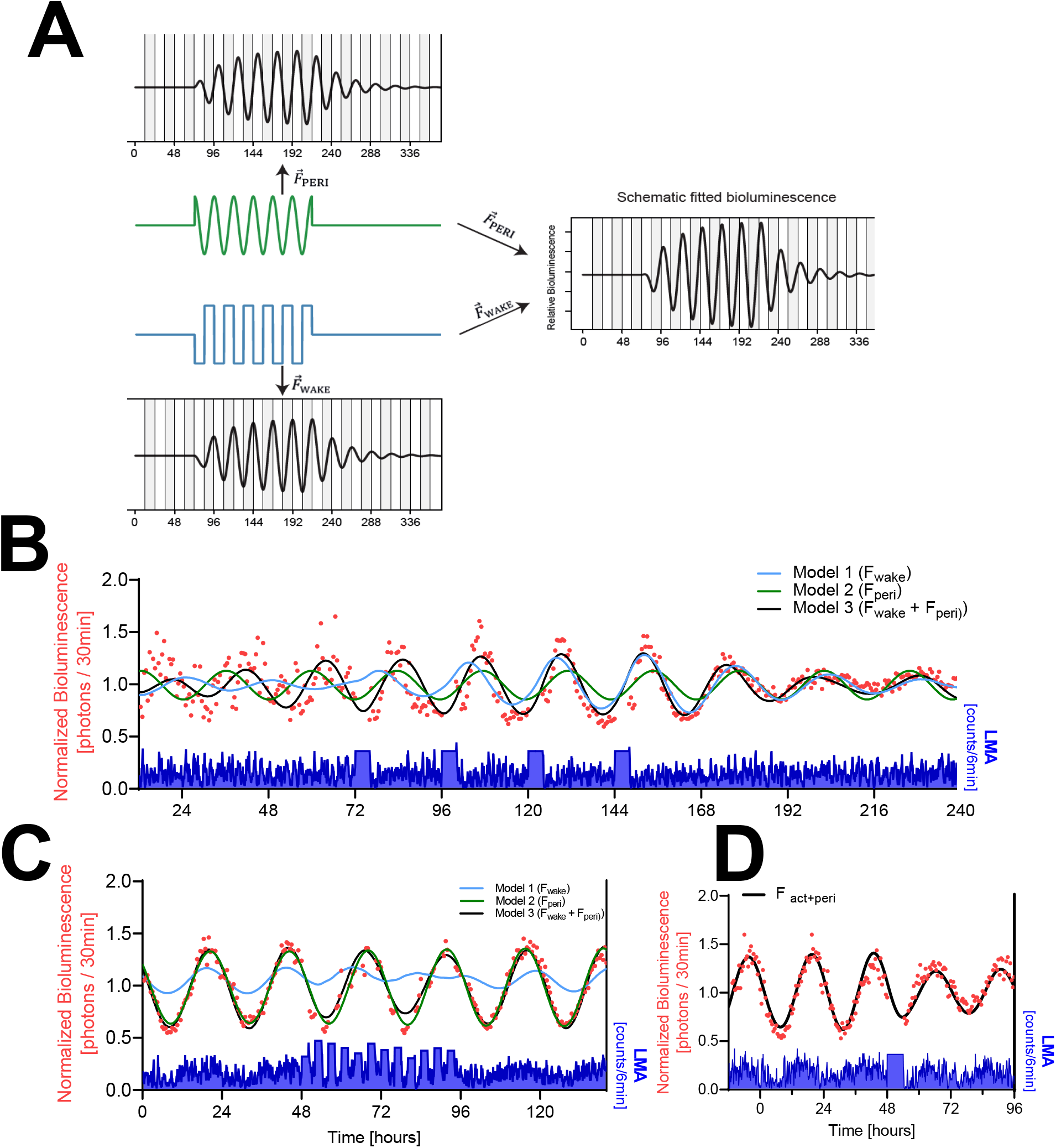
Mathematical modeling of PER2 bioluminescence dynamics. (**A**) Schematic view of a driven damped harmonic oscillator. The oscillation (black) is assumed to be driven by two forces: 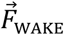 (blue) and 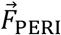 (green). In our model 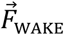 is based on LMA (here simplified as a square wave) and 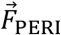 on a sinewave. Left panels show the individual effect of each force on the oscillator. Both forces start at t=72 h and end at t=216 h, illustrating the waxing and waning of the resulting rhythm amplitude. Right panel shows the resulting changes in PER2 bioluminescence when combining both forces. Note that combining the two forces increased amplitude and changed the phase of the oscillation. In this example, the amplitude of the peripheral circadian force is flat at beginning and end to illustrate that the oscillation is not self-sustained. (**B**) Modeling of the SCNx experiment with the full model (Model 3; black line) driving oscillations in PER2-bioluminescence or with either 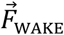 (Model 1; blue line) or 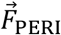 (Model 2; green line) as the only driving force. Red symbols; 30-minute PER2-biolumininescence averages. Lower graph in blue: 6-minute LMA values. (**C**) Simulation of the PER2-biolumininescence in the 2hOnOff experiment using parameter estimates listed in Table 1. Details as in B. (**D**) Simulation of peripheral PER2-biolumininescence in the 6h SD experiment, using parameter estimates obtained in C (see Table 1).

**Table 1:** Parameter estimates obtained in the model optimization in the SCNx and 2hOnOff experiments. Damping constant, natural angular frequency, and the 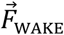 coefficient were optimized in the SCNx experiment and then used to predict the results of the 2hOnOff experiment. Amplitude and phase of 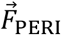 were optimized for both experiments separately. The natural angular frequency defines the intrinsic period of the harmonic oscillation with 0.288 rad * h^−1^ corresponding to a period of 21.8h and its square, 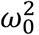 is referred to as the string constant. Phase of the sine function describing 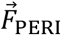 is expressed as radians and corresponds to maximum values reached at time 11.8 and 19.4h for the SCNx and 2hOnOff experiment, respectively. Values in parenthesis represent the 95%-CI.

| Parameters | SCNx | 2hOnOff |
| --- | --- | --- |
| $\gamma$ (damping constant) | 0.0155 (0.0103 - 0.0223) [$\text{h}^{-1}$] | |
| $\omega_0$ (natural angular frequency) | 0.288 (0.285 - 0.291) [$\text{rad} \cdot \text{h}^{-1}$] | |
| $\beta$ (coefficient $\vec{F}_{\text{WAKE}}$) | 6.81e-5 (4.48e-5 – 9.01e-5) | |
| Model Intercept | 0.92 (0.90-0.95) | 0.91 (0.89 - 0.92) |
| $A$ (amplitude $\vec{F}_{\text{PERI}}$) | 2.92e-3 (2.58e-3 - 3.32e-3) | 3.87e-3 (3.80e-3 - 4.00e-3) |
| $\varphi$ (phase $\vec{F}_{\text{PERI}}$) | 4.73 (4.6 - 4.85) | 2.78 (2.78 - 2.82) |

We first optimized the parameters of the model describing PER2-bioluminescence dynamics in the SCNx experiment (data from Fig. 2), and then predicted PER2-bioluminescence under the 2hOnOff experiment although 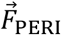 ‘s amplitude and phase again required optimization as both differed between the two experiments (non-overlapping 95%-CI for both in Table 1). With the parameters listed in Table 1, the model captured the dynamic changes in PER2 bioluminescence in the SCNx experiment with high precision including the residual rhythmicity in baseline, the reinstated pronounced rhythmicity during the 4 SDs, and its subsequent dampening thereafter (Fig. 4B, black line, Table 2). The model also accurately captured average bioluminescence dynamics in the 2hOnOff experiment (Fig. 4C, black line, Table 2). It furthermore predicted an 18% reduction of PER2 bioluminescence amplitude during the 2hOnOff protocol compared to baseline (relative amplitudes estimated by the model: 0.31 versus 0.38 [a.u.]; Fig. 4C), consistent with our hypotheses and the 27% reduction in PER2 bioluminescence amplitude observed in the 4 mice in which amplitude did decrease (see Fig. 3C). Finally, we applied the model to predict the peripheral PER2-bioluminescence data obtained in the 6h SD experiment (Fig. 1A) using the parameters of the intact mice in the 2hOnOff experiment (Table 1). Also the results of this experiment could be accurately predicted with the model, including the higher PER2-bioluminescence levels reached over the initial recovery hours after SD and the subsequent longer-term reduction in amplitude (Fig. 1A and 4D, Table 2). The model could not reliably predict the central PER2-bioluminescence dynamics mainly due to an earlier phase of the central compared to the peripheral signal (Suppl. Fig. 7D). Moreover, the increase in PER2 levels immediately following the SD (Fig. 1A) was missed by the model, further underscoring the tissue-specific relationship between time-spent-awake and PER2 requiring the model to be optimized according to the tissue under study.

**Table 2:**
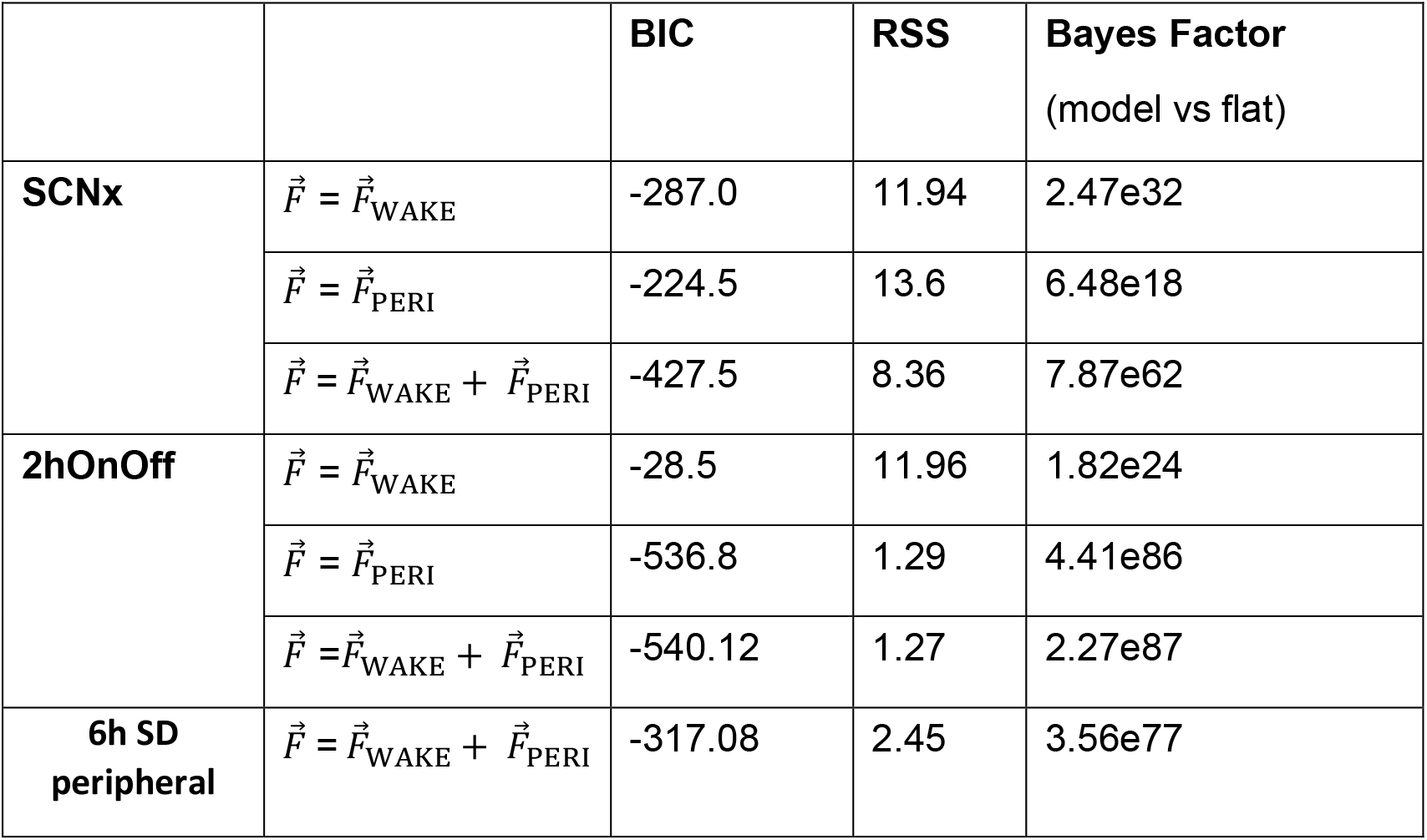
Fitting statistics for the SCNx and 2hOnOff experiments using a single force (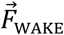 or 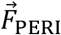) or 2 forces combined (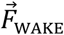 and 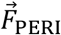). Bayesian Information Criterion (BIC) for each model (lower is better; a BIC difference between two competing models larger than 10 is considered strong support for the model with the lower value). Residual sum of squares (RSS) minimized by the model (lower values reflect a better fit). Support of the driven harmonic model compared to a flat model using Bayes Factor (>100 is considered a ‘decisive’ support for the driven harmonic model). Values should be compared only within the same experiment because variance and sample size differed among the experiments. Fit statistics for the prediction of the peripheral 6h SD results using the parameters optimized for the 2hOnOff experiment with the full model (Table 1) have been included for completeness.

The model accurately captured peripheral PER2 bioluminescence dynamics under three different experimental conditions. The model’s robust performance prompted us to assess *in silico* whether 1) both forces are required to predict PER2 dynamics, 2) lesioning the SCN affects the forces exerted on PER2 under undisturbed, DD conditions, and 3) differences in the circadian phase could predict the opposing response in PER2 amplitude among mice in the 2hOnOff experiment.

To evaluate if 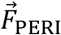 and 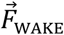 combined are required to predict PER2 dynamics, we compared the performance of the model by dropping these two modeling terms; i.e., removing either 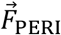 or 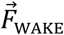. Although without 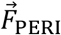 the model is unable to capture the residual PER2 rhythmicity in the baseline of the SCNx experiment (Fig. 4B, ‘model 1’ until 72h), removing 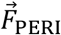 from the model did not change the coefficient obtained for 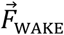 (8.25e-5 which is still within the 95%-CI estimated in the full model; see Table 1). However, this simpler model (only 4 free parameters to estimate compared to 6 in the full model) could not reliably predict the bioluminescence data of the SCNx and 2hOnOff experiments (Fig. 4B and -C; blue lines) and fit statistics indicated a poorer fit (i.e., a higher BIC score, see Table 2). Moreover, without 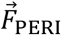, it took longer for the PER2 bioluminescence to reach high amplitude levels compared to the full model, because of 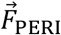 being in phase with the timing of the SDs in the full model. With the same strategy we evaluated the model’s performance when dropping 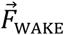 (Fig. 4B and -C; green lines). As the sleep-wake distribution no longer affects the model, 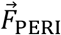 dynamics are unperturbed throughout the experiment and can therefore be solved as a sine-wave in the differential equation. Fit statistics for the SCNx experiment were poor compared to the full model as the effects of SDs on PER2 amplitude could not be captured (Fig. 4B, Table 2). Removing 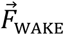 when modeling the 2hOnOff experiment resulted in BIC scores similar to those obtained with the full model (Table 2), although the amplitude reduction during the 2hOnOff protocol could not be captured (Fig. 4C). Together with the modelling of the SCNx and the 6h SD experiments, these results demonstrate that both forces are needed to predict PER2 dynamics under the tested experimental conditions.

To answer the second question, we compared the forces driving PER2 bioluminescence in the baseline recordings of the 2hOnOff (intact mice) and the SCNx experiments. The model estimated that the average absolute force (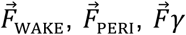, and 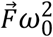 combined) exerted on the relative PER2-bioluminescense levels was more than twice as high in intact mice compared to SCNx mice (2.11e-2 vs. 0.90e-2 h-2). This large difference was not due to a difference in 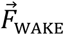 which was somewhat lower in intact mice (2hOnOff vs. SCNx; 2.01e-3 vs. 2.22e-3 h-2). Although 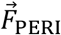 was higher by 32% (2.45e-3 vs. 1.85e-3 h-2) this difference did not substantially contribute to the higher amplitude of the PER2 rhythm in intact mice. To illustrate this, we substituted the value of 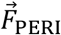 obtained in the 2hOnOff baseline with the lower value obtained in the SCNx baseline, which resulted in a 17% amplitude reduction of the PER2 rhythm. The presence of a circadian sleep-wake distribution impacted PER2 amplitude to a larger extent: running the simulation with the parameters estimated for the 2hOnOff experiment but with the sleep-wake distribution of the SCNx mice resulted in a 32% reduction. Therefore, the circadian sleep-wake organization is an important contributor to high amplitude oscillations in PER2. In the 2hOnOff experiment, the larger PER2 momentum resulted in a 4 times higher string and damping forces (2hOnOff vs. SCNx; 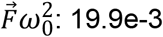 vs. 4.4e-3 h-2; 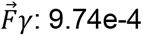 vs. 2.52e-4 h-2) that together with the larger 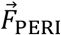 underlie the larger average absolute force. Moreover, this analysis demonstrated that the direct effects of 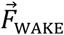 and 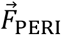 on PER2 bioluminescence do not depend on an intact SCN and that the magnitude of the two forces are comparable.

Although the model predicted the expected decrease in PER2-bioluminescense rhythm amplitude in the 2hOnOff experiment, this decrease was not observed in the mean bioluminescence data. The individual data showed, however, that in 4 mice this intervention did decrease PER2-bioluminescence amplitude, while in the remaining 2 amplitude increased (Fig. 3C). With the model we addressed the third question: Do circadian phase differences predict the opposite effects of the 2hOnOff intervention on amplitude? Surprisingly, when systematically varying the phase of 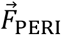 during the baseline prior to the 2hOnOff intervention (Suppl. Fig. 7C), we found that at phase advances larger than 2.5h, the model predicted an increase of PER2 amplitude during the subsequent 2hOnOff protocol instead of a decrease (illustrated in Suppl. Fig. 7A for a 6h phase advance). Circadian phase differences among animals might relate to individual differences in period length of their free-running rhythms that accumulate over time. We did not find evidence for a difference in PER2 bioluminescence or LMA phase at the end of the baseline recording (2.5 days under DD) between animals that showed a decrease in PER2-bioluminescence amplitude compared to animals that showed an increase. However, during the first day of recovery following the 2hOnOff protocol (5.5 days under DD), we observed a ca. 2h phase advance in bioluminescence and 1h phase advance in LMA in the 2 mice that increased their amplitude during the preceding 2hOnOff protocol (Suppl. Fig. 7B, left), compared to the 4 mice for which we observed the anticipated decrease in PER2 amplitude (Suppl. Fig. 7B, right). Whether these differences in phase contributed to the opposite response cannot be answered with the current data set. Nevertheless, the model yielded a perhaps counterintuitive but testable hypothesis by demonstrating that phase angle between the sleep-wake distribution and peripheral circadian clock-gene rhythms is an important variable in predicting outcome and emphasizes the importance of carefully controlling the initial conditions.

## Discussion

In this study we assessed the contribution of the sleep-wake distribution to circadian peripheral PER2 rhythmicity. We presented three key findings, supporting the notion that sleep-wake state is indeed an important factor in determining the circadian amplitude of peripheral changes in PER2: i) sustained bouts of waking and sleep were associated with increased and decreased PER2 bioluminescence, respectively; ii) repeated SDs temporarily reinstated robust rhythmicity in PER2 bioluminescence in behaviorally arrhythmic mice, and iii) mathematical modeling suggest that PER2 dynamics is best understood as an harmonic oscillator driven by two forces: a sleep-waking dependent force 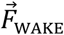 and a SCN-independent and sleep-wake independent, circadian peripheral force 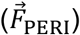.

### HOW DOES WAKEFULNESS INCREASE PER2?

*Per2* transcription can be initiated from its cognate E-boxes by CLOCK/NPAS2:ARNTL. This transcriptional activation is at the core of the TTFL and drives the circadian changes in PER2. Enforced wakefulness not only affects *Per2* levels but also modulates the expression of other components of the TTLF (Mang & Franken, 2015; Hor et al. 2019). The SD-evoked increase in *Per2* expression could therefore be mediated through other clock genes in the circuitry, as was demonstrated by the differential SD-evoked response in *Per2* levels in mice lacking the core clock genes *Npas2* and both *Cry1* and *-2* genes (Franken et al., 2006; Wisor et al., 2002). Apart from a TTFL-mediated activation, *Per2* transcription can be induced by other signaling molecules directly acting on elements within the *Per2* promotor (Schibler et al., 2015). For example, ligand-bound glucocorticoid receptors can induce *Per2* transcription by binding to their glucocorticoid response elements (Cheon et al., 2013; So et al., 2009). Similarly, cAMP response element (CRE)-binding protein (CREB), heat-shock factor 1 (HSF1), and serum-response factor (SRF) can directly activate *Per2* transcription through CREs, heat-shock elements, and CArG-boxes, respectively, present in the *Per2* gene (Gerber et al., 2013; Camille Saini et al., 2012; Tamaru et al., 2011; Travnickova-Bendova et al., 2002). Through these pathways, *Per2* responds to stress, light, temperature, blood-borne systemic cues, and cellular activation as an immediate early gene (IEG). Because of this, *Per2* can appear rhythmic even in the absence of a functional TTFL, provided these signaling pathways fluctuate cyclically (Kornmann et al., 2007). In behaviorally arrhythmic SCNx animals the residual PER2 rhythms we observed might similarly result from SCN-independent corticosterone (Andrews, 1968) or body temperature (Satinoff & Prosser, 1988) rhythms, or, alternatively, might be TTFL-driven locally (Sinturel et al., 2021).

Important for the current study is that several of the pathways known to directly influence *Per2* expression are activated by either spontaneous and/or enforced waking [e.g., corticosterone (Mongrain et al., 2010), temperature (Hoekstra et al., 2019), *Hsf1* and *Srf* (Hor et al., 2019), pCREB (Cirelli & Tononi, 2000)] and are therefore good candidates linking sleep-wake state to changes in PER2. The observed changes in PER2-bioluminescence were rapid and suggest that increases in protein can occur within an hour of spontaneous wakefulness. Other studies document that PER genes can indeed be rapidly transcribed and translated. For instance, a light pulse given at CT14 leads within an hour to a significant increase in *Per2* transcript in the SCN (Yan & Silver, 2002). One study reported a large increase in hepatic *Per2* transcript levels within 1h after food presentation in fasted rats (Wu et al., 2010), underscoring the ability of this transcript to rapidly adapt to homeostatic need. *Per2* translation is not solely dependent on *de novo* transcription and, for example in the SCN, light was shown to promote *Per2* translation (R. Cao et al., 2015), suggesting that transcription would not be necessary to increase PER2 protein levels. Such mechanism could also underlie the very fast (<30min) 1.5-fold increase in PER2 protein observed in fibroblasts after serum shock (R. Cao et al., 2015).

Finally, as the PER2 protein levels measured are the net result of translation and degradation also sleep-wake dependent changes in PER2-degradation rate may contribute both to its increase during wakefulness and its decrease during sleep. PER2 degradation is crucial in setting TTFL period and the timing and stability of the circadian sleep-wake distribution (Chong et al., 2012; D’Alessandro et al., 2017). One established pathway leading to PER2 degradation involves *Casein kinase 1* (*Ck1*) mediated phosphorylation (Eide et al., 2005) followed by the recruitment of the ubiquitin ligase *β-transducin repeat-containing proteins* (*Btrc)* (Masuda et al., 2020; Ohsaki et al., 2008; Reischl et al., 2007). Others kinases, such as *Salt-inducible kinase 3* (*Sik3*) (Hayasaka et al., 2017) and phosphorylation-independent ubiquitin ligases, such as *Transformed mouse 3T3 cell double minute 2* (*Mdm2*) (Liu et al., 2018, p. 2), also target PER2 for degradation. Using a modelling approach to estimate the role of *Btrc* in circadian period length, Reischl (Reischl et al., 2007) estimated a linear decay rate of 0.18/h for PER2 degradation, which is not inconsistent with the approximately 0.10/h net decay rate we observed for the PER2-bioluminescene during sleep. However, the dynamics of PER2 degradation have been assessed in a circadian context exclusively, and effects of sleep-wake state have not been quantified previously.

The obvious next step is to determine which pathway(s) contributes to the wake-driven changes in PER2 protein. We already established that the sleep-deprivation incurred increase in *Per2* in the forebrain partly depended on glucocorticoids (Mongrain et al., 2010). Along those lines, restoration of daily glucocorticoid rhythms in adrenalectomized rats reinstates PER2 rhythms in several extra-SCN brain areas (Segall & Amir, 2010). To determine the contribution of the aforementioned wake-driven factors, a genetic screen could be deployed where one-by-one the regulatory elements in the *Per2* promoter are mutated, and the effect of sleep-wake driven *Per2* changes is assessed. This approach has already been taken for the GRE and CRE elements in the *Per2* promoter to test their respective roles in circadian phase resetting and integrating light information (Cheon et al., 2013; So et al., 2009; Travnickova-Bendova et al., 2002).

### INSIGHTS FROM THE MODEL

The model accurately captured the main features of peripheral PER2 dynamics observed in all three experiments, thereby giving further credence to the notion that sleep-wake state importantly contributes to the changes in PER2 observed in the periphery. Moreover, it demonstrated that the large amplitude of the circadian PER2 rhythm in intact mice is likely the result of the momentum gained in the harmonic oscillator through the daily recurring sleep-wake distribution. This is conceptually different from a currently accepted scenario, in which direct and indirect outputs from the SCN assure phase coherence of locally generated self-sustained circadian rhythms (Schibler et al., 2015). According to this model, loss of amplitude observed at the tissue level in SCNx mice is caused by phase dispersion of the continuing rhythms in individual cells. Among the SCN outputs thought to convey phase coherence are feeding, locomotor activity and changes in temperature. Because these outputs all require or are associated with the animal being awake, it can be argued that in both models the circadian sleep-wake distribution is key in keeping peripheral organs rhythmic.

The harmonic oscillator model further showed that, although important, the sleep-wake force alone was not sufficient to predict PER2 dynamics. In addition to account for the residual PER2 rhythms observed in undisturbed SCNx mice, the SCN-independent and sleep-wake-independent circadian force greatly improved the performance of the model in both experiments. The synergistic effect of both forces (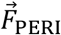 and 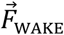) was needed to explain the rapid response to the SDs observed in the SCNx experiment and also in maintaining robust circadian rhythms during the SDs in the 2hOnOff experiment as illustrated in Fig. 4B and -C, respectively. Furthermore, this synergistic effect greatly depended on the relative phase of the two forces as we could illustrate *in silico* for the 2hOnOff experiment: a relative subtle change in the phase of 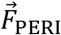 might underlie the increase (instead of the predicted decrease) in PER2 amplitude in 2 of the 6 mice recorded.

Which pathways set the phase of 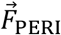 and whether it is truly independent of 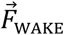, as assumed in the model, our current results cannot answer. In an earlier modeling effort using a similar approach in SCN-intact, light-dark entrained mice, we also required a second force to correctly predict the phase of the observed rhythm in brain *Per2* expression (Curie et al., 2013). In that publication we based the second force on the pattern of corticosterone production sharply peaking at ZT11 just prior to the light-dark transition. The phase of 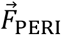 in the 2hOnOff experiment, which followed a more gradual, sine-wave function of which values became positive shortly after ZT11 (extrapolated from the preceding LD cycle), seems consistent with this. In the SCNx experiment, the phase of 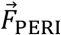 was positioned ~7h earlier with a positive driving force starting at the end of each of the SDs. As SD is accompanied by an increase in corticosterone (Mongrain et al., 2010), the phase of 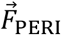 could be associated with corticosterone signaling also in the SCNx experiment. Thus in intact mice, the SCN output would dictate the phase of corticosterone production in the adrenals (and thus that of 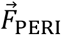), while in SCNx mice the phase of the corticosterone rhythm can be reset by stressors such as SD. As PER2 and *Per2* levels in the SCN seem insensitive to SD (Curie et al., 2015; Zhang et al., 2016), this could explain why the phase of 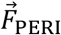 is maintained in sleep-deprived SCN-intact mice. Moreover, *ex vivo* experiments demonstrated that the adrenal gland can generate bona-fide circadian rhythms in corticosterone release independent of the SCN (Andrews, 1968; Engeland et al., 2018; Kofuji et al., 2016), even though SCNx is generally thought to abolish rhythms in circulating corticosterone levels (Moore & Eichler, 1972). While rhythmic corticosterone release represents a plausible candidate contributing to 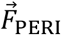, especially considering its role in synchronizing peripheral clocks (Balsalobre et al., 2000; Cuesta et al., 2015; Dickmeis, 2009; Le Minh et al., 2001), our current results cannot rule out other sources underlying or contributing to 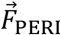. Above we argued that also wakefulness (i.e., 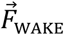 in the model) could influence peripheral PER2 through corticosterone signaling, further complicating the issue. Indeed, adrenalectomy was found to reduce (but not abolish) the SD-induced increase in *Per2* expression in the forebrain (Mongrain et al., 2010). However, enforced but not spontaneous wakefulness is accompanied by increases in corticosterone and the model could predict PER2 dynamics without having to distinguish between the two types of waking. Therefore, other candidate signals among those listed above must be considered to understand the biological basis of 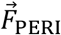 and 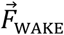.

Model optimization yielded an unexpected short 21.8h period for the natural frequency of the PER2 oscillator, which, in addition, differed from the 23.7h period we set for 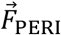. While in the intact mice of the 2hOnOff experiment it is difficult to independently determine 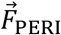 ’s period, we estimated a 23.7h period length for the residual PER2 rhythmicity observed during baseline in SCNx mice, which we assume is driven by 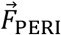. In intact mice kept under constant conditions, SCN output drives behavioral sleep-wake rhythms and synchronizes peripheral clock-gene rhythms forcing the entire system to oscillate at the intrinsic period of the SCN. Similarly, in behaviorally arrhythmic SCNx mice, we assume that the period of the observed residual PER2 rhythm reflects that of the only remaining driver, 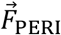, as 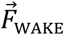 is no longer rhythmic and direct effects of the SCN are absent. *Ex vivo* experiments showed that periods vary among tissues and do not depend on whether tissues were obtained from an intact or SCNx mouse (Cederroth et al., 2019; Yoo et al., 2004), pointing to tissue-specific TTFLs, which we assume to underlie the intrinsic rhythmicity of both 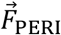 and PER2 bioluminescence in our experiments. The difference in period length of 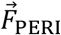 and that of the intrinsic PER2 oscillator therefore suggests that 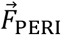 is not of renal origin; i.e. the tissue which contributed most to the bioluminescence signal we recorded.

### TISSUE SPECIFICITY OF THE RELATIONSHIP BETWEEN SLEEP-WAKE STATE AND PER2

Our data demonstrated that both central and peripheral PER2-bioluminescence dynamics are affected by sleep-wake state, not only after SD but also after spontaneous periods of wakefulness. Despite this general observation, we found clear tissue-specific differences: the 6h SD elicited an immediate increase in the central PER2 signal, while in the periphery this increase occurred several hours later, confirming our earlier findings in brain versus liver and kidney (Curie et al. 2015). Tissue specificity was also observed after spontaneous bouts of wakefulness: in the brain, PER2 bioluminescence immediately increased after the animal woke up and continued to do so until the end of the waking bout, whereas in the kidney the increase in PER2 bioluminescence became apparent only 5-10 min after the transition. Similar differences in PER2 dynamics were observed after falling asleep, albeit in opposite direction. Also the model suggested a tissue-specificity, as central dynamics of the 6h SD experiment could not be accurately predicted with the parameters optimized on peripheral PER2 bioluminescence data. In its current form, the model describes the global effects of external forces (the sleep-wake distribution and the SCN independent circadian force) on the behavior of the oscillator, but not their acute effects. Translated into molecular terms, the model only makes predictions on the TTFL aspect of PER2 regulation, not on PER2 as an IEG. Accordingly, the model cannot capture the fast changes at the transitions. Similarly, the short-lasting high levels of PER2 observed in the brain immediately after the 6h SD might reflect an IEG response rather than the state of the TTFL oscillator which would explain why the model could not predict it. As sleep-wake states are brain states, it stands to reason that changes in brain PER2 levels capture more of the acute IEG effects than in the periphery. One could test this hypothesis by quantifying the PER2 response after activating a peripheral tissue provided this can be achieved without affecting sleep-wake state as well.

### DO CHANGES IN BIOLUMINESCENCE REFLECT CHANGES IN PER2 LEVELS?

The method we implemented to quantify PER2 protein levels presents advantages over previous methods used. It enabled us to acquire data at a time resolution needed to link changes in PER2 to sleep-wake state transitions in individual mice. Moreover, because of the within-subject experimental design, there is a substantial reduction in data variability and, as illustrated with the effects of individual phase on PER2 amplitude in the 2hOnOff experiment, we could assess the presence of individual differences in the initial conditions that might influence experimental outcome. Finally, the number of mice needed for these experiments has been greatly reduced while obtaining better quality data.

A limitation of this method is the assumption that changes in bioluminescence reflect changes in PER2 protein levels. Using western blot, we previously validated that changes in bioluminescence during baseline and after a 6h SD indeed reflect changes in PER2 protein (Curie et al., 2015). Nevertheless, substrate availability can importantly contribute to the signal as demonstrated in the experiment in which we delivered luciferin in the drinking water. Even the use of osmotic mini-pumps does not guarantee constant delivery, as its release rate is temperature dependent (Alzet, manufacturer’s notes) and bioluminescence’s increase during wakefulness might therefore result from the accompanying increase in temperature during this state. Arguments against a possible temperature effect on bioluminescence changes comes from a study in which, using the same osmotic mini-pumps, the expression of two clock genes known to oscillate in anti-phase could be confirmed (Ono et al., 2015), which would not be possible if temperature was the main determinant of bioluminescence. In the current data, the circadian rhythm in wakefulness in the CAG mice was not accompanied by changes in bioluminescence and if anything decreased during the active phase. Moreover, we observed that the circadian rhythms of subcutaneous temperature and bioluminescence are ca. 4h out of phase (data not shown), supporting that the large circadian changes in bioluminescence are not driven by changes in luciferin availability. Nevertheless, to fully exclude that potential rate-limiting availability of luciferin contributed to the fast sleep-wake evoked changes in PER2, we would require confirmation with other techniques sufficiently sensitive to quantify the small but consistent protein changes during sleep and wake.

## CONCLUSIONS

In this study, we used a unique combination of methods allowing us to collect high-resolution data of sleep-wake state in conjunction with PER2 levels and found that the sleep-wake distribution profoundly affects PER2 bioluminescence both short- and long-term. Such behavior-dependent plasticity of the time-keeping machinery in tissues peripheral to the SCN enables the organism to respond to challenges as time restricted feeding experiments have demonstrated (Damiola et al., 2000; Saini et al., 2013). Besides its importance in regulating feeding and energy homeostasis (Bass & Takahashi, 2010), the clock circuitry also plays a prominent role in sleep homeostasis (Franken 2013). PER2 seems perfectly suited as an integrator of sleep-wake state and circadian time, because it is sensitive to a variety of sleep-wake driven signals. Our model suggests that having a large amplitude rhythm protects from acute disturbances of sleep as observed in the 2hOnOff experiment, while sleep depriving arrhythmic SCNx mice had immediate and large effects on PER2. These rapid effects could only be achieved through the synergistic effect of a second force that we found to be independent of the SCN and the sleep-wake distribution. The coordination of the sleep-wake force and this second force in the model was critical in predicting the effects of sleep-wake interventions on PER2. Research on the nature of this second force would therefore be important to facilitate phase resetting and normalize disrupted clock gene rhythms under conditions of jet lag and shift work, complementing strategies aimed at altering the timing of the central pacemaker.

## Acknowledgements

We are greatly indebted to all who helped with the sleep deprivations: Lisa Härri, Charlotte Hor, and Jeffrey Hubbard, and especially to those who sacrificed their sleep during the graveyard shifts: Kostas Kompotis, Simone Mumbauer, Violeta Castelo-Szekely and Sonia Jimenez. We also thank Sonia for her help with the sleep-wake annotation of EEG/EMG files.

## Material and Methods

### MICE AND HOUSING CONDITIONS

To measure peripheral PER2 bioluminescence levels we made use of the *Per2::Luc* KI construct (Yoo et al., 2004). The *Per2::Luc* KI construct was originally generated on a C57BL/6J-129 mixed background and subsequently brought onto a C57BL/6J (B6) background by backcrossing for at least 11 generations. These mice where then crossed with outbred SKH1 mice (Crl:SKH1-Hrhr; Charles River) to create hairless *Per2::Luc* KI mice. We used male homozygous *Per2::Luc* KI B6 and hairless heterozygous *Per2::Luc* KI SKH1-B6 hybrid (here referred to as SKH1 mice) mice. Mice were kept under a 12 h-light/12 h-dark cycle with light- and dark-onset referred to as Zeitgeber time (ZT)-0 and −12, respectively. Age at time of recording varied between 12 and 24 weeks. Food and water was available *ad libitum,* and after surgery mice were singly housed. All experiments were approved by the Ethical Committee of the State of Vaud Veterinary Office Switzerland under license VD2743 and 3201.

### SOURCE OF BIOLUMINESCENCE AND LUCIFERIN’S ROUTE OF ADMINISTRATION

Because *Per2::Luc* KI mice express ubiquitously luciferase and the RT-Biolumicorder cannot discriminate between different sources of bioluminescence, we assessed which peripheral organ(s) was/were the major source of bioluminescence. Two male heterozygous *Per2::Luc* SKH1 mice were implanted with an osmotic mini-pump (model 1002; 35mg/mL luciferin) and five days later lightly anaesthetized with 2.5% isoflurane and imaged for 60 seconds (Xenogen IVIS Lumina II) around ZT6. The main source of dorsal bioluminescence overlapped with the expected location of the kidney, whereas ventrally almost no bioluminescence was detected (see Suppl. Fig. 3A). Most bioluminescence quantified during the experiment is of dorsal origin due to the orientation of the mouse relative to the PMT, suggesting that the kidneys are the main source of peripheral bioluminescence in *Per2::Luc* SKH1 mice.

In a second pilot experiment, we investigated the optimal route of luciferin administration. Although luciferin administration via drinking water has been used before to measure bioluminescence (Saini et al., 2013, Iwano et al., 2018, Hall et al., 2018, Sinturel et al. 2021), we were concerned that this route could limit luciferin availability in a circadian fashion because drinking behavior has a strong circadian rhythm (Bainier, Mateo, Felder-Schmittbuhl, & Mendoza, 2017). To address these concerns, we made use of mice expressing constitutively luciferase under control of the synthetic CAG promotor [(Y. A. Cao et al., 2004), Jackson catalog number 008450], thus allowing for testing of circadian fluctuating levels of luciferin. Mice received luciferin via the drinking water or via an osmotic mini-pump and served as their own control. Four male CAG mice were housed for two subsequent experiments in constant darkness in the RT-Biolumicorder. During the first experiment, 0.5 mg/mL luciferin was dissolved in the drinking water. At the end of this experiment, mice received subcutaneously an osmotic mini-pump (Alzet, model 1002) under light anesthesia (isoflurane; 2-4% mixed with O_2_) containing 70 mg/mL of luciferin and could recover for two days, before bioluminescence and activity was monitored for the second experiment in the RT-Biolumicorder.

### SURGICAL PROCEDURES AND EXPERIMENTAL DESIGN

Experimental design of the 3 main experiments has been depicted in Suppl. Fig. 1 with Panel A illustrating the central and peripheral recordings of PER2 bioluminescence alongside EEG/EMG and LMA before, during, and after a 6h SD, and Panels B and C the SCNx and the 2hOnOff experiments, respectively. In the latter 2 experiments peripheral PER2 bioluminescence and LMA were recorded.

#### Sleep-wake state determination in parallel with PER2 bioluminescence

Mice were implanted with electroencephalogram (EEG) and electromyogram (EMG) electrodes under deep ketamine/xylazine anesthesia. Three gold-plated screws (frontal, parietal and cerebellar) were screwed into the skull over the right cerebral hemisphere. Two additional screws were used as anchor screws. For the EMG, a gold wire was inserted into the neck musculature along the back of the skull. For brain delivery of D-luciferin, a cannula (Brain Infusion Kit1, Alzet) was introduced stereotaxically into the right lateral ventricle (1 mm lateral, mm posterior to bregma and 2.2 mm deep) under deep anesthesia (ketamine/xylazine; intraperitoneally, 75 and 10 mg/kg, respectively), and connected to the mini pump. A depression (diameter 2 mm) was made in (but not through) the skull in a region of the left frontal cortex (approximate coordinates 2 mm lateral to midline, 2 mm anterior to bregma), in which a glass cylinder (length 4.0 mm; diameter 2.0 mm) was positioned and fixed with dental cement. The EMG and three EEG electrodes were subsequently soldered to a connector and cemented to the skull. The cerebellar screw served as a reference for the parietal and frontal screw and the EMG. After the first recovery day, mice were habituated to the weight of the wireless EEG recording system by attaching a dummy of same size and weight to their connector. Two days before habituation to the RT-Biolumicorder, mice were implanted with the osmotic mini-pump (model 1002, Alzet; luciferin 35mg/mL) under light anesthesia. 8-10 days post-surgery, mice were placed in the RT-Biolumicorder at the end of the light phase (~ZT10-ZT12) for two days in LD to habituate to the novel environment. At the end of the second habituation day, the dummy was replaced with a wireless EEG (Neurologger, TSE Systems GmbH). After two-and-a-half days of baseline recording in constant darkness, mice were sleep deprived for six hours at a time they were expected to rest (ZT0 under LD conditions) by gentle handling as described (Mang & Franken, 2012). In short, mice are left undisturbed as long as they do not show signs of sleep. Sleep is prevented by introducing and removing paper tissue, changing the litter, bringing a pipet in the animal’s proximity, or gentle tapping of the cage. As opposed to what the term might suggest, mice are not handled. After SD, mice were placed back into the RT-Biolumicorder for the subsequent two recovery days.

#### SCNx experiment

Four SKH1 mice were recorded over the course of the experiment and served as their own control. Briefly, their PER2 bioluminescence rhythm was monitored before SCNx (once), under undisturbed conditions post-SCNx (twice), and after the second measure under SCNx conditions, the mice were subjected to the repeated 4h SDs.

Bilateral lesion of the two SCNs was performed stereotaxically (Kopf Instruments, 963LS, Miami Lakes, FL, USA) under ketamine/xylazine anesthesia (intraperitoneal injection, 75 and 10 mg/kg, at a volume of 8 mL/kg). Two electrodes (0.3 mm in diameter) were introduced bilaterally at the following coordinates (position of the frontal electrode: anteroposterior using bregma as reference: ±0.2mm lateral, +0.5mm bregma, depth: −5.9mm; the second electrode was positioned 0.7 mm posterior to the frontal one). Electrolytic lesions (1 mA, 5 sec) were made using a direct current (DC) lesion device (3500, Ugo Basile, Comerio, Italy). After lesion, mice were housed in constant dark (DD) conditions for at least 10 days to verify absence of circadian organization of overt behavior. Activity was quantified using passive infrared sensors (Visonic SPY 4/ RTE-A, Riverside, CA, USA). ClockLab software (Actimetrics, Wilmette, IL, USA) was used for data acquisition and analyses.

#### Surgeries for tethered EEG/EMG recordings

SKH1 mice (n=8) were implanted with EEG and EMG electrodes as described previously (Mang & Franken, 2012) to determine sleep-wake state. The surgery took place under deep xylazine/ ketamine anesthesia. Briefly, six gold-plated screws (diameter 1.1 mm) were screwed bilaterally into the skull over the frontal and parietal cortices. Two screws served as EEG electrodes and the remaining four screws anchored the electrode connector assembly. As EMG electrodes, two gold wires were inserted into the neck musculature. The EEG and EMG electrodes were soldered to a connector and cemented to the skull. Mice recovered from surgery during several days before they were connected to the recording cables in their home cage for habituation to the cable and their environment, which was at least 6 days prior to the experiment. The habituation to the room and the recovery from the two-day SD-procedure took place under LD 12:12 conditions.

During the baseline recording and sleep deprivation days, red light at very low intensity was present to allow the experimenters to observe the mice. Mice were sleep deprived for two hours according to the ‘gentle handling’ method.

#### Mice for bioluminescence data collection

SKH1 mice (n=6) were implanted with an osmotic mini-pump (Alzet, 1002, luciferin concentration: 35 mg/mL; blue flow moderator) two days before the habituation. At the end of the light phase (~ZT10-ZT12), mice were moved from their cage to the RT-Biolumicorder for 2-3 days of habituation in LD. They were housed for 2.5 days in DD, after which the 2hOnOff protocol was initiated at light onset (ZT0) under the preceding LD conditions. At the start of each SD, mice were moved from the RT-Biolumicorder and placed into a novel cage that was in the same room as the EEG-implanted mice. Fifteen minutes before the end of each SD, mice were brought back to their RT-Biolumicorder cage.

### DATA COLLECTION OF SLEEP-WAKE STATE

#### Simultaneous recording of sleep-wake state and PER2 bioluminescence

Batteries (Hearing Aid; Ansmann, 312 PR41, 1.45 V 180 mAh) were inserted into the Neurologger. This insertion was timed with the clock of the computer that controlled the RT-Biolumicorder to *post hoc* align the EEG/EMG signals with the bioluminescence signal. In addition, time stamps provided by the SyncBox (NeuroLogger, TSE) were used to verify the start and end time of the EEG/EMG recording. The cerebellar electrode was used as a reference for both EMG and EEG. Data was sampled at 256 Hz. The frontal signal was subtracted from the parietal signal (EDF Browser) to support sleep-wake state determination by enhancing the identification of slow waves and theta waves within the same trace. The data was subsequently loaded in Somnologica (Somnologica 3, MedCare) to determine offline the mouse’s behavior as wakefulness, REM sleep or NREM sleep per 4-second epochs based on the EEG and EMG signals. Wakefulness was characterized by EEG activity of mixed frequency and low amplitude, and present but variable muscle tone. NREM sleep (NREM) was defined by synchronous activity in the delta frequency (1-4 Hz) and low and stable muscle tone. REM sleep (REM) was characterized by regular theta oscillations (6-9 Hz) and EMG muscle atonia.

#### 2hOnOff experiment

EEG and EMG signals were recorded continuously for 96 h. The recording started at the beginning of the subjective rest phase, ZT0 of the preceding LD cycle. The analog EEG and EMG signals were amplified (2,000×) and digitized at 2 kHz and subsequently down sampled to 200 Hz and stored. Like the EEG and EMG traces obtained with the Neurologger, the data was imported in Somnologica and sleep-wake state was determined per 4-second epochs. LMA was monitored with passive infrared activity (ActiMetrics, US, Wilmette) and recorded with ClockLab (ActiMetrics, US, Wilmette).

### DATA ANALYSIS

#### Route of luciferin administration

Circadian time was determined to inspect the circadian changes in bioluminescence relative to locomotor activity. To this end, the period length per mouse was determined based on activity measurements (1-min resolution) by chi-square analysis in ClockLab. Subsequently, the activity and bioluminescence data were folded according to the period. The activity data was averaged per 10 minutes and activity onset was visually determined for each mouse and set at CT12. The aligned activity and bioluminescence data were subsequently averaged per circadian hour.

#### Spontaneous sleep-wake state and bioluminescence

Bioluminescence and activity were sampled at a resolution of 4-seconds, which is the same resolution of the epochs for sleep-wake state determination. Data processing was subsequently performed in MatLab 2017b (The MathWorks, Inc., Natick, Massachusetts, United States) and R (version 4.0.0). Linear trends were removed from the signal by the build in function *detrend* in Matlab and R (*pracma* package). Subsequently, the bioluminescence signal was expressed relative to the overall mean per mouse to account for inter-individual differences. Changes in PER2 would be expected to occur at a slower rate then 4-seconds. Therefore, sleep-wake and bioluminescence data are averaged per blocks of 3 minutes.

For the sleep-wake transition analysis, 3 min intervals in which the mouse was awake for more than 50% (i.e. 23 or more 4s epochs) were deemed awake otherwise as asleep. Based on this new 3-min sleep-wake sequence clear sleep-to-wake and wake-to-sleep transitions were selected according to the following criteria: at least 9 min (three 3-min intervals) of the initial state had to be followed by at least 15 min (five 3-min intervals) of the other state. Transitions were followed both forward and backwards in time as long as state did not change. Bioluminescence for all 3-min intervals of a transition were expressed as a percentage of the level reached at the transition; i.e., the average between the level reached in the last 3 min prior to the transition and the first 3min after the transition. Transitions were aligned according to time of the state transition (time 0) and then averaged first within and then across mice. Average time courses were reported for the longest time spent in state after the transition to which all mice contributed.

### 2hOnOff experiment

Bioluminescence data obtained five minutes before and ten minutes after the sleep deprivations was excluded from analysis. Subsequent data normalization of the bioluminescence data was as above. To determine the influence of the 2hOnOff protocol on the strength of the ongoing circadian oscillation of the circadian distribution of sleep and wake, as well on PER2 bioluminescence, an estimation of amplitude by sinewave fitting was done (MatLab, 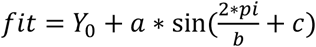).

### Modeling PER2 bioluminescence with a damped harmonic oscillator

The temporal dynamic of PER2 bioluminescence was modeled according to the equation of motion describing a driven damped harmonic oscillator:

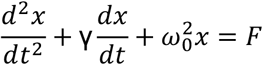

where x is the displacement of the oscillator, γ is the damping constant of the model, and 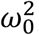 the string constant defining the natural frequency of the model. The driving forces used in this model are the locomotor activity (LMA) representing waking 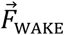, and a circadian force 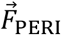, represented as a sinewave. The momentary force exerted on the oscillator is represented as the sum of these two forces (see Fig. 4A):

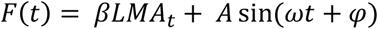

Where *β*, *A* and *φ* are respectively the coefficient applied on locomotor activity amount, the amplitude of the circadian sinewave force, and the phase of the circadian sinewave force. These coefficients were the free parameters to be optimized in the model. *ω* is the angular velocity of the sinewave and was set to 2*pi/23.7, based on the residual PER2-bioluminescence rhythm present in the baseline of SCNx mice. To solve this equation and optimize for parameters that best describe the observed PER2 bioluminescence, we proceeded as follows: The relative bioluminescence data from both experiments was averaged across mice using 30-minute bins. LMA was averaged across mice using 6-minute bins. Locomotor activity could not be measured directly during sleep deprivations (SD) and was estimated by assessing the increase in LMA during SDs measured in EEG-implanted mice, which was found to be 2.4 times higher compared to average baseline levels. Thus, the SD effect was estimated using 2.4 times the mean activity observed during baseline (i.e. 181.9 and 180.2 for the SCNx and 2hOnOff, respectively).

To solve the 2^nd^ order ordinary differential equation (ODE) of the driven harmonic oscillator, we transformed it into the following system of two first order ODEs describing the change of position and speed of our oscillator:

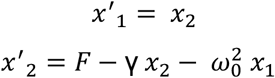

We then used the 4^th^ order Runge-Kutta (RK4) numerical method to approximate the solution using a fixed time step of 0.1 hour. In the SCNx experiment, initial values of speed (*x*_2_(0)) and position (*x*_1_(0)) were set to 0. For the 2hOnOff experiment the model was generated for 20 days prior experiment to reach a steady-state using replication of LMA observed in baseline (T0-T24). The position and speed of the oscillator at the end of the 20 days were taken as initial values for the fitting. We optimized the model for the following parameters: intercept (equilibrium position of the oscillator), natural frequency, damping constant, and coefficient for the force exerted by LMA, amplitude of circadian force and phase of circadian force. We optimized the fitting minimizing the residual sum of squares (RSS) between predicted position of our model and the observed PER2 bioluminescence level. The box-constrained PORT routines method (nlminb) implemented in the optimx/R package (Nash & Varadhan, 2011) was used to minimize the RSS.

The goodness of fit of the model was assessed as follows. We assumed that the model errors follow a normal distribution and computed a Bayesian Information Criterion (BIC) value for the model according to:

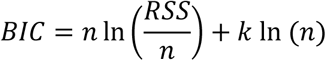

Where n is the number of observations, and k is the number of parameters of the model. We approximated the Bayes Factor (BF) between our model and a flat model (linear model with intercept only) as follow:

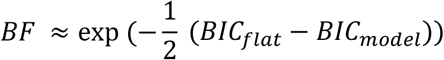

To compute confidence interval of our model parameters we used 500 Monte-Carlo simulations and calculated a confidence interval for our parameters based on 95% empirical quantiles (95%-CI). The method was adapted from the code of Marc Lavielle [Inria Saclay (Xpop) and Ecole Polytechnique (CMAP), Université Paris-Saclay, France] for nonlinear models, available here: http://sia.webpopix.org/nonlinearRegression.html

### Statistics

Statistics were performed in R (version 4.0.0), SAS (version 9.4), and Prism (version 7.0), with the threshold of significance set at α=0.05. Performed statistical tests are mentioned in the text and figure legends of the result section.

## Supplementary Figures

**Supplementary Figure 1:**
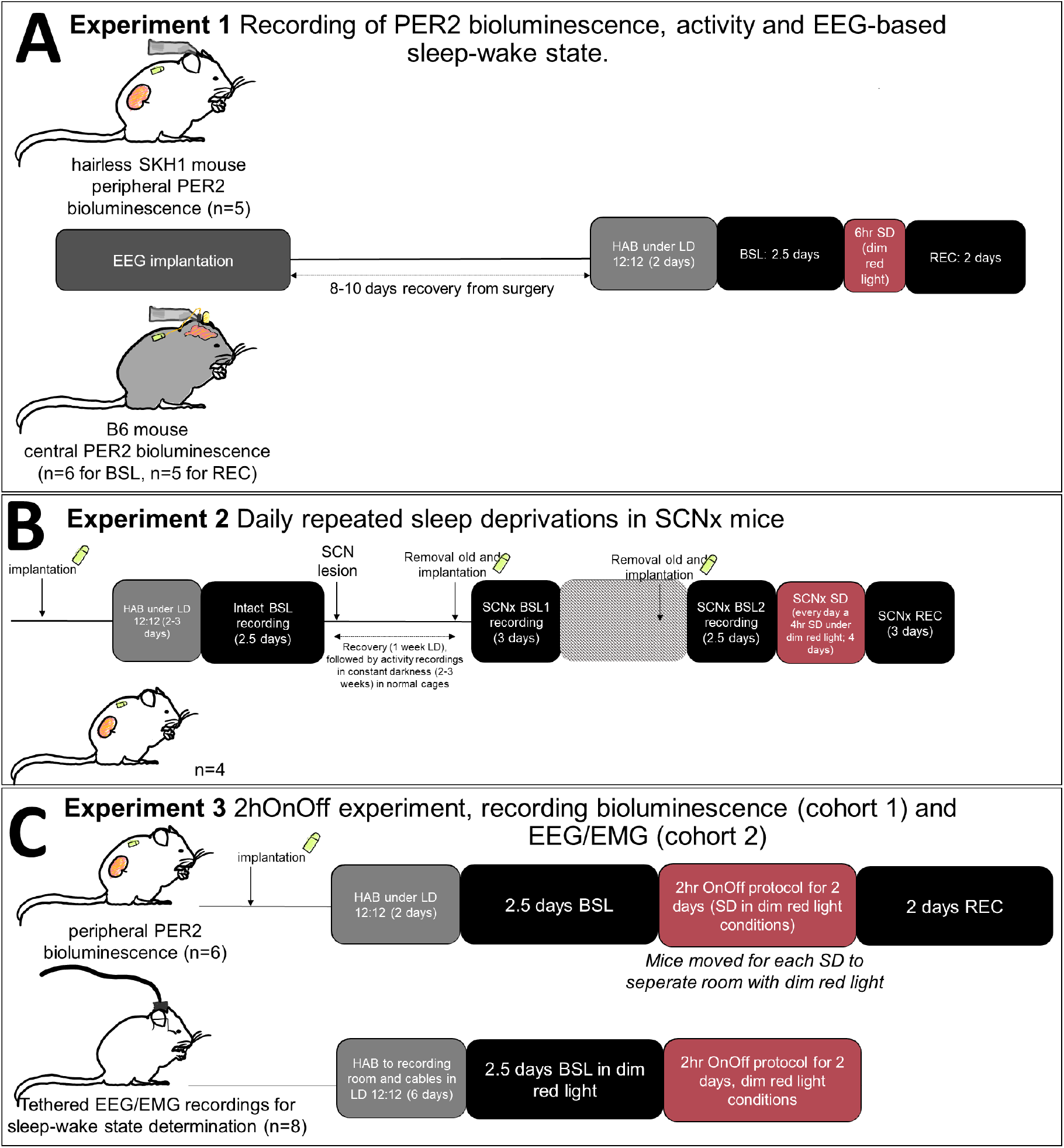
Experimental protocol for the three experiments. HAB: habituation; BSL: baseline; SD: sleep deprivation; REC: recovery; SCNx: SCN lesion.

**Supplementary Figure 2:**
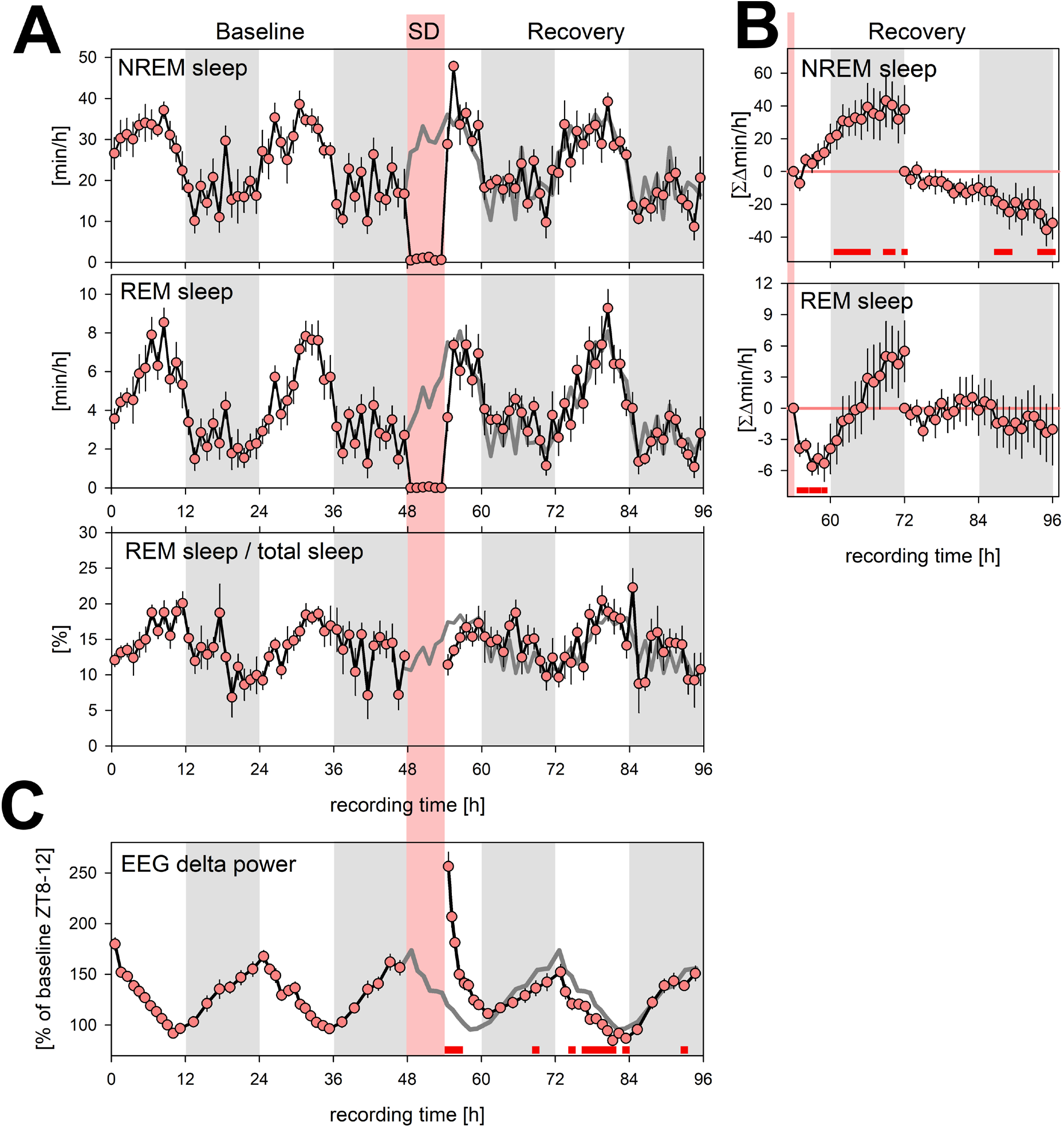
Sleep phenotyping of SKH1 mice. Four-day time course of time spent asleep and EEG delta power (1-4 Hz) during baseline, 6h sleep deprivation (SD; ZT0-6, t=48-54h; pink area), and recovery in SKH1 mice (n=8) under LD12:12 (grey areas denote the 12h D-periods) conditions. (**A**) NREM sleep (top) and REM sleep (middle), as mean minutes over hourly intervals (±1 SEM). Also depicted is REM sleep as % of total sleep (NREM + REM sleep; bottom panel). Dark-grey line during SD and recovery represent baseline time course (mean over the 2 baseline days) for comparison. (**B**) Accumulated differences (min/h) from baseline separately for recovery day 1 (t=54-72h) and day 2 (t=72-96h). During recovery day 1 mice accrue ca. +38.8±14.6 min extra NREM sleep, which is again lost during recovery day 2 (−31.5±9.6 min). Mice lose −5.3±1.7 min of REM sleep during first 5 recovery hours after SD. This additional deficit is offset during the remaining of recovery day 1 resulting in a non-significant increase of 5.5±2.9 min. (**C**) Mean EEG delta power during NREM sleep (±1 SEM) calculated for 12 and 6 percentiles during the L- and D-periods, respectively (except for L-period recovery day 1; 8 percentiles) to which an equal number of 4s-epochs scored as NREM sleep contributed (per LD period and mouse). Delta power is considered an EEG correlate of sleep need decreasing when sleep prevails (L-period) and increasing when animals are mostly awake (D-periods). SD results in high immediate values that quickly revert to baseline. Red squares underneath curves represent hourly intervals in which values differed from baseline, post-hoc paired t-tests p<0.05.

**Supplementary Figure 3:**
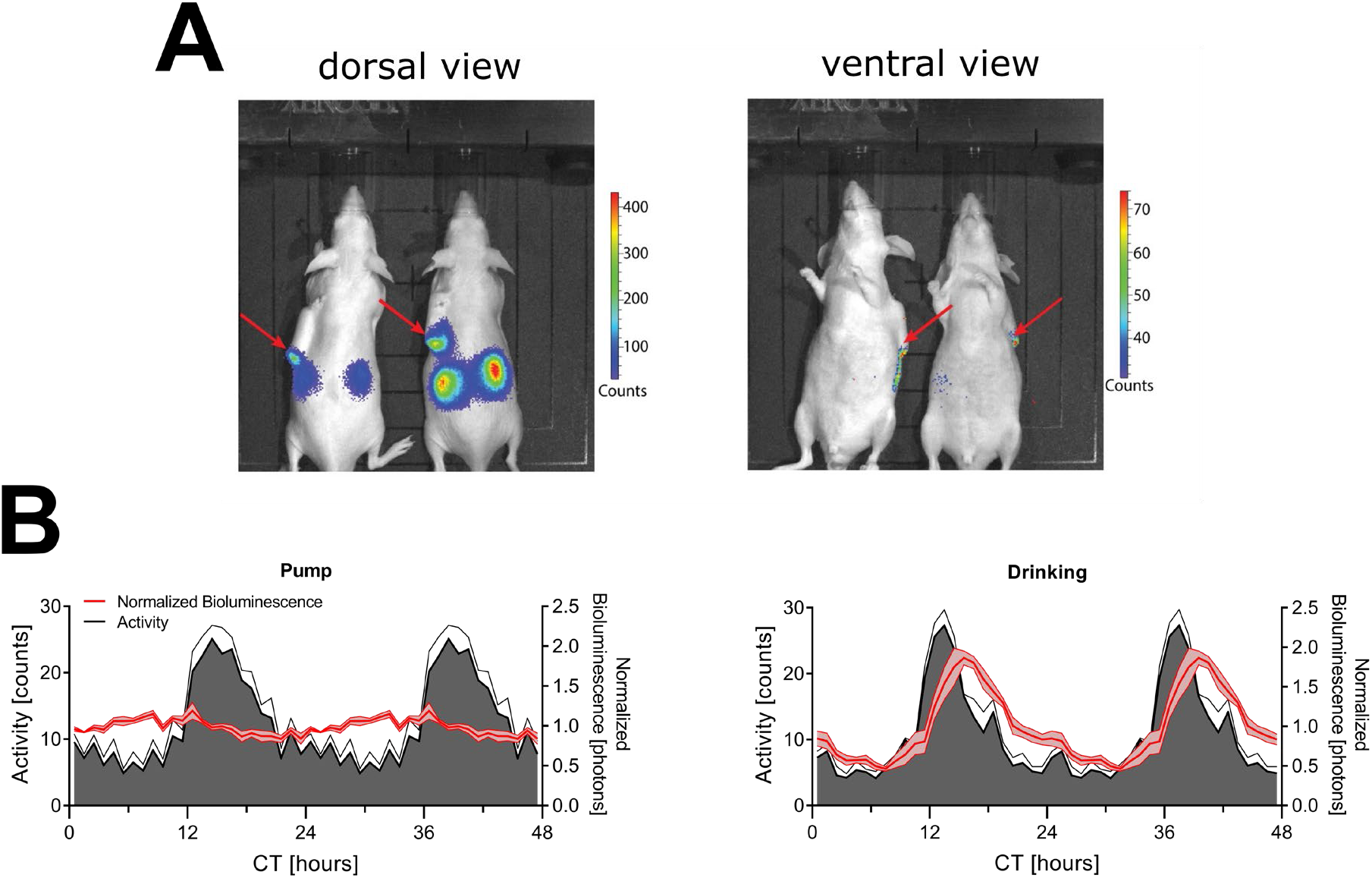
Sources of bioluminescence and different luciferin administration routes. (**A**) Dorsal (left) and ventral (right) imaging of mice in IVIS Xenogen system. Note that the scale bar of the dorsal view has higher values than the scale bar of the ventral view. Images are taken between ZT6-8. The red arrows indicate the location of the osmotic minipump flow moderator. (**B**) Mice constitutively expressing luciferase under the control of the synthetic CAG promoter received luciferin via an osmotic mini-pump (left) or in their drinking water (right) (n=4). Graphs are double plotted to aid visualization. CT = circadian time.

**Supplementary Figure 4:**
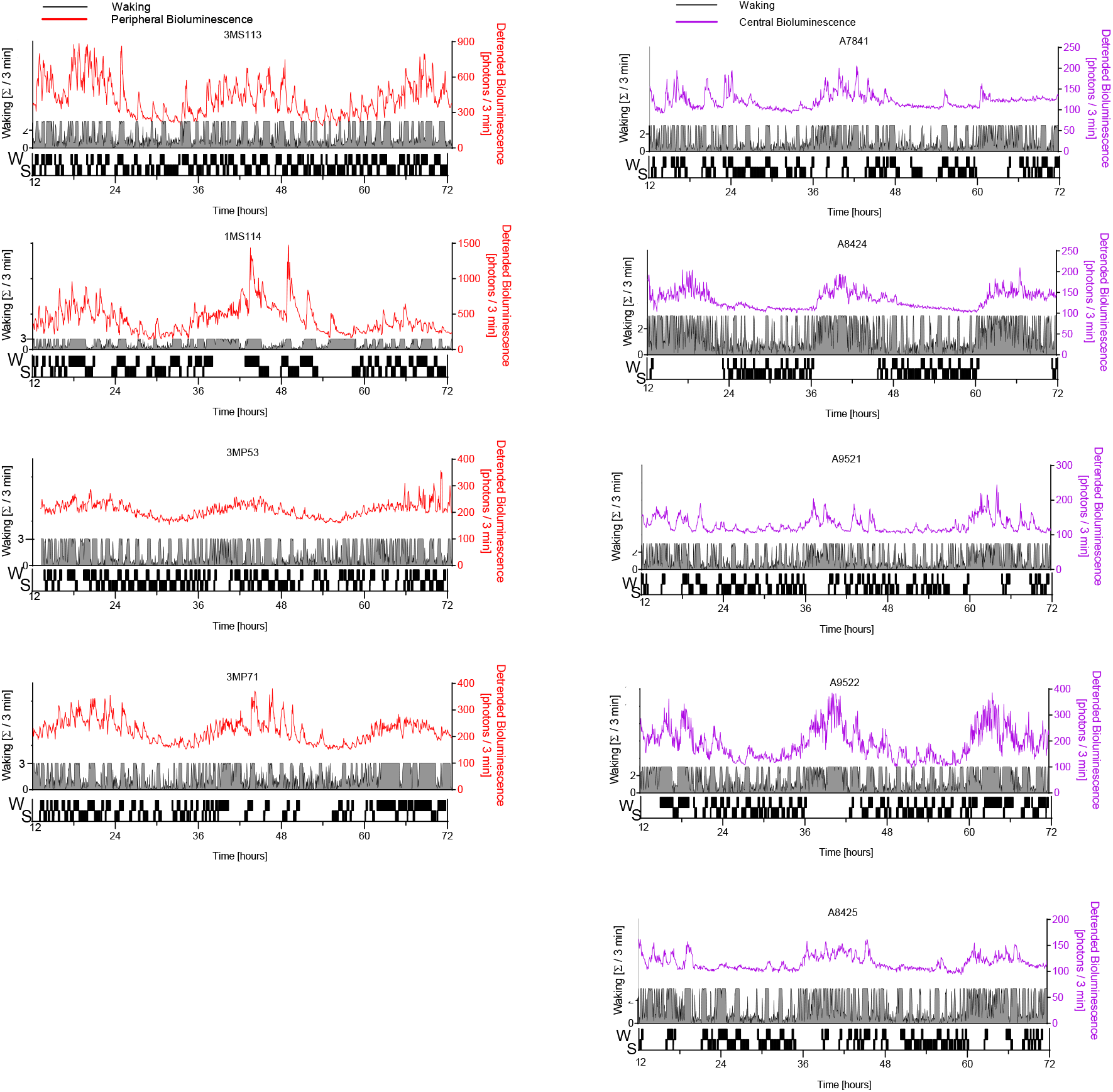
(with Fig. 1B) Bioluminescence co-detected with EEG-based sleep-wake state in the other mice. Left, peripheral bioluminescence (red traces); right central bioluminescence (magenta traces). Legend as in Fig. 1B.

**Supplementary Figure 5:**
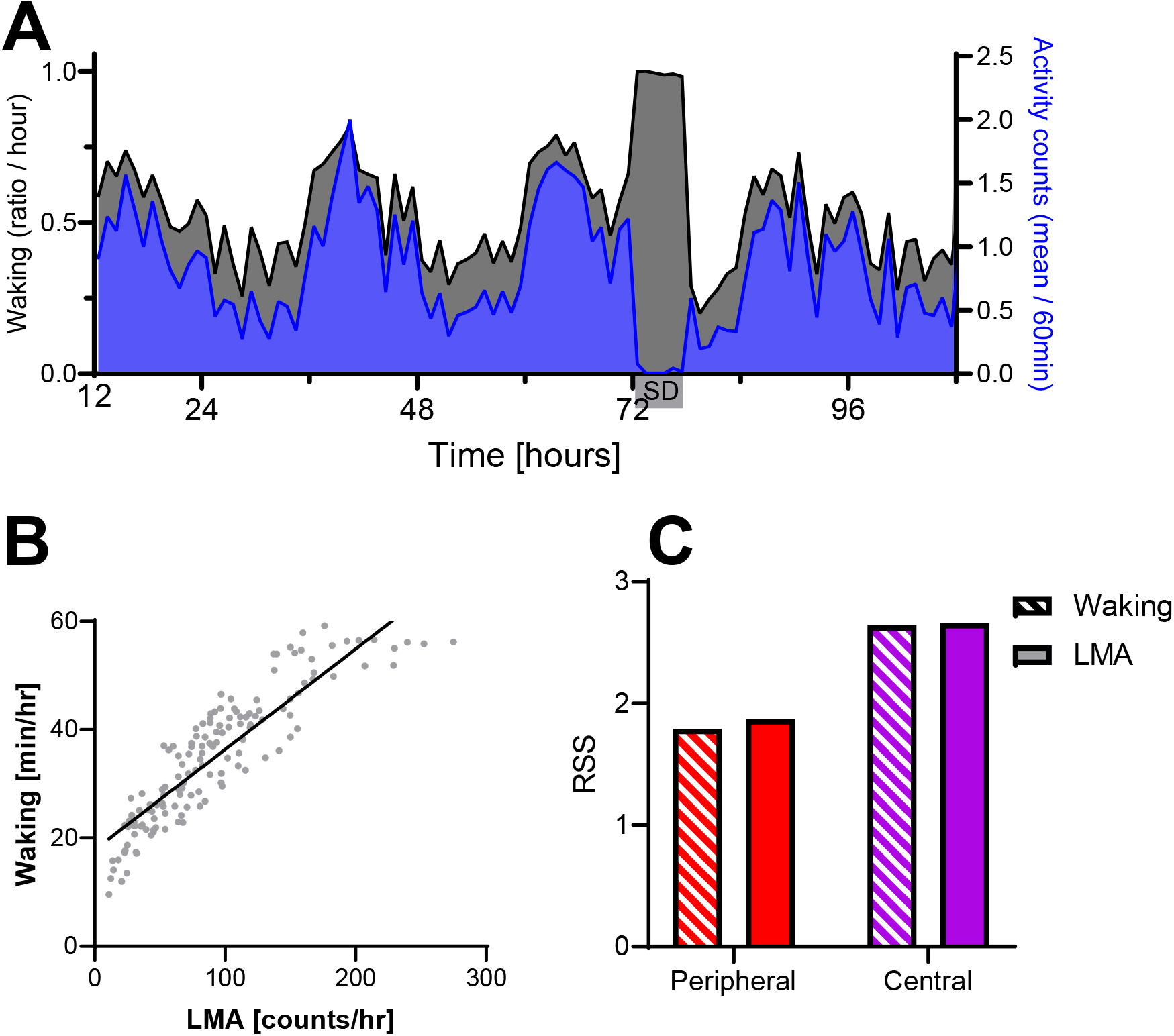
(with Fig. 4) Locomotor activity (LMA) as proxy for wakefulness (**A**)EEG-based waking (black) and locomotor activity (blue) from 5 mice used to determine central PER2 bioluminescence during a two-and-a-half day baseline recording (12-72h), a sleep deprivation (72-78h) and the recovery days. Note that no activity was measured during the SD as mice were not in the cage with the activity sensor at that time. (**B**) LMA and waking measured in the 2hOnOff protocol EEG experiment, linear regression, p<0.0001, R^2^=0.79. (**C**) Similar residual sum of squares (RSS) for the prediction of PER2 dynamics were obtained when using waking (striped) or LMA (filled) as an input for 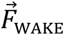 in central (left) and peripheral (right) recordings.

**Supplementary Figure 6:**
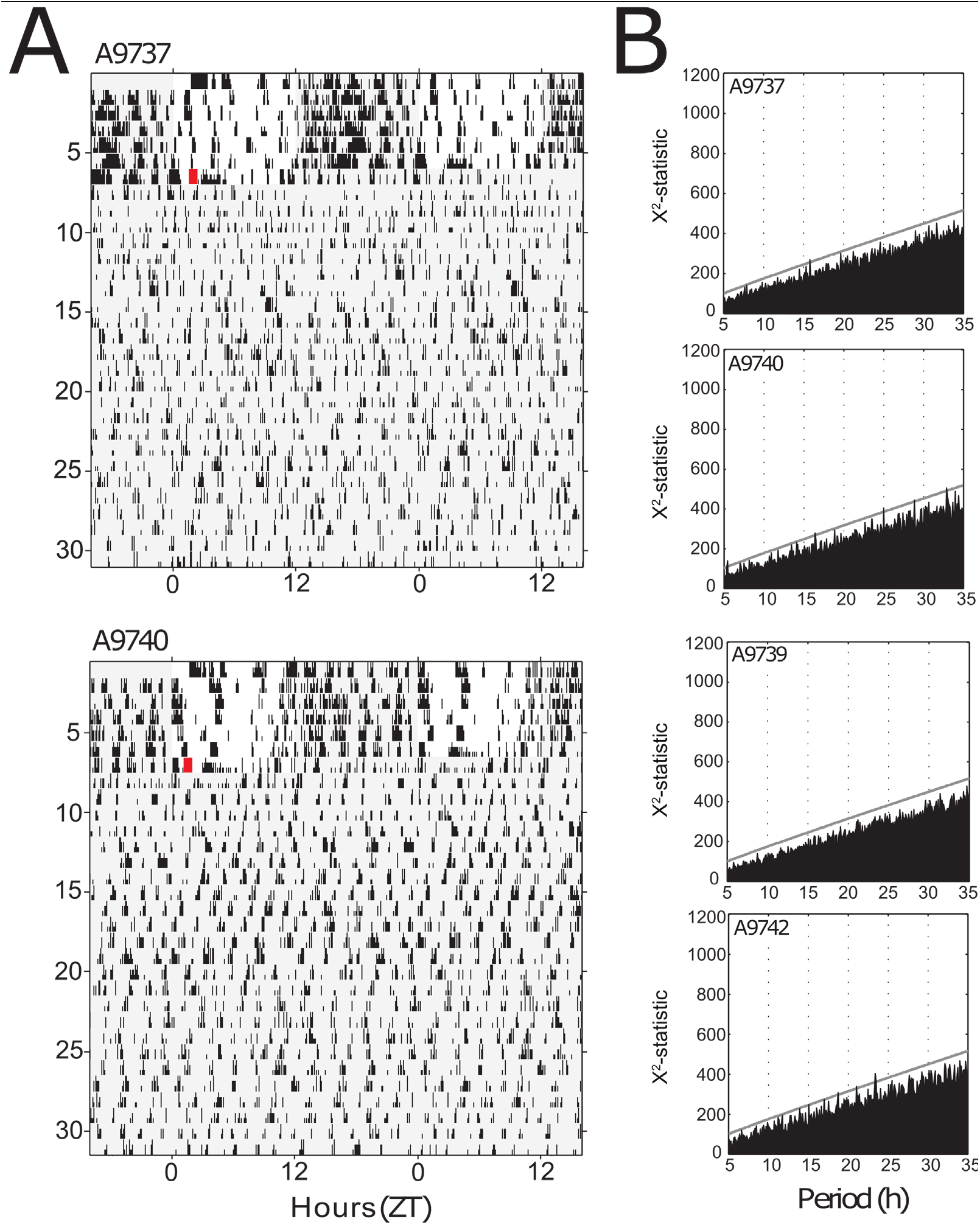
SCN-lesion eliminates circadian organization of locomotor activity. (**A**) Examples of double-plotted actograms for two mice kept under LD12:12 condition (days 1-7 vertical axis) before bilateral lesion of the SCN (red square) and subsequently released under DD conditions (days 7-31). Light-grey areas denote darkness. (**B**) Chi-square periodograms of locomotor activity under DD following SCN lesion. Analysis in upper two graphs concern the mice illustrated in A. Grey lines in each panel indicate P=0.001 significance level.

**Supplementary Figure 7:**
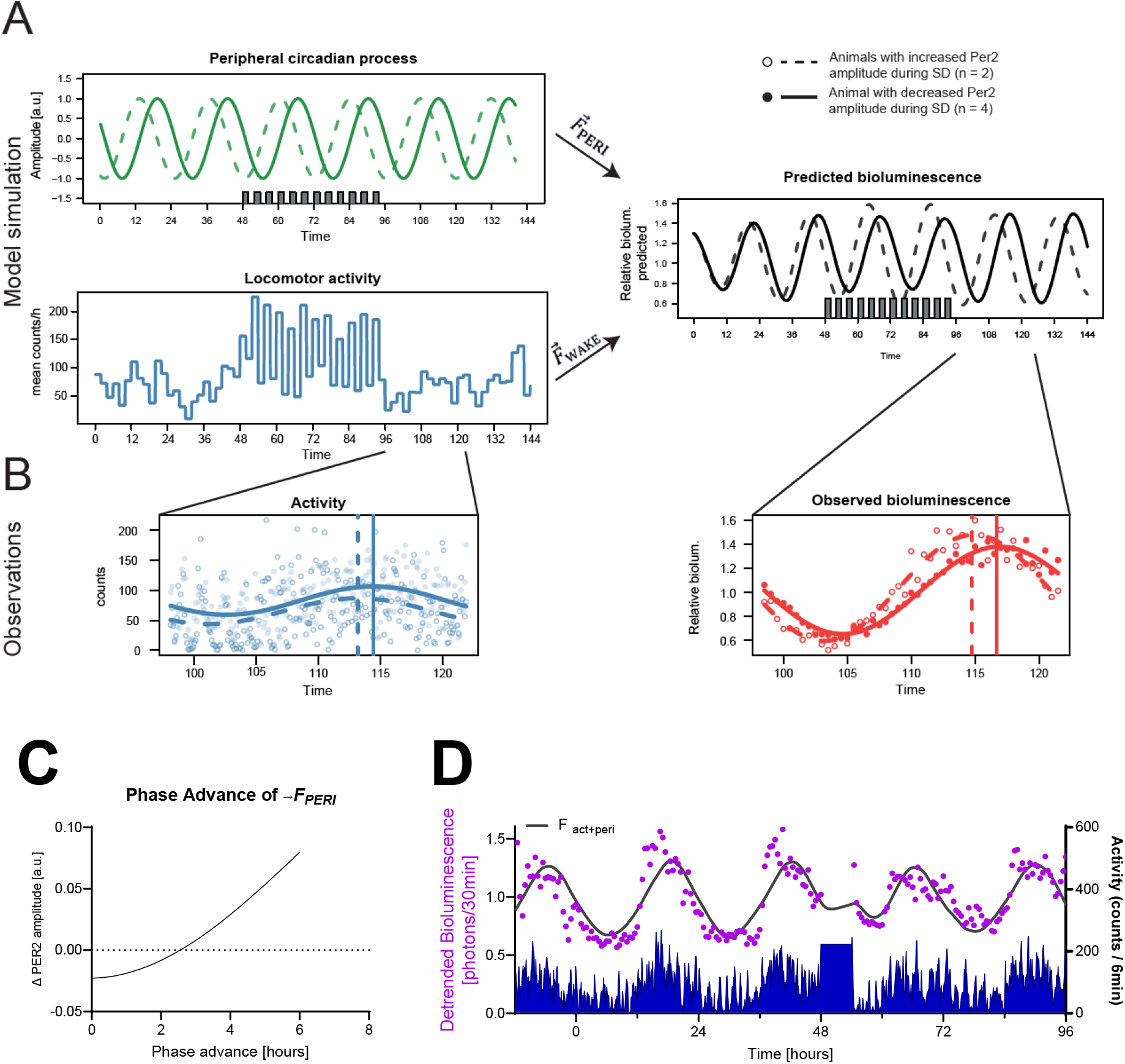
Predicting PER2 bioluminescence considering 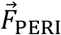 ‘s phase (A-C) and bioluminescence emitted by cortical regions (D). (**A**) Model simulation using fitted parameters as in Fig. 4C (solid lines) and assuming a 6h phase advance of 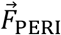 (dashed lines). Amplitude of predicted PER2 level (black lines) increased during 2hOnOff when phase was advanced (dashed), while it decreased using the fitted parameters (solid line). 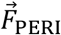 in green; 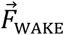 in blue. (**B**) Phase advance observed for PER2 bioluminescence (red) and wake (blue) in mice at the end of sleep deprivation for mice which increased PER2 bioluminescence amplitude during 2hOnOff (empty circles, n=2) vs those in which amplitude decreased (filled circles, n=4). Phase (vertical line) was calculated using sine-wave fitting (solid and dashed lines). (**C**) Predicting the effect of a phase advance of 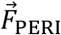 on PER2 amplitude reveals that when the phase advance > 2.5h, PER2 amplitude will increase. (**D**) PER2 bioluminescence predicted based on the parameters obtained for peripheral bioluminescence are not able to predict the earlier, sharp increase of PER2 bioluminescence at activity onset, nor the acute increase following sleep deprivation.

